# The distribution of plasmid fitness effects explains plasmid persistence in bacterial communities

**DOI:** 10.1101/2020.08.01.230672

**Authors:** Aida Alonso-del Valle, Ricardo León-Sampedro, Jerónimo Rodríguez-Beltrán, Javier DelaFuente, Marta Hernández-García, Patricia Ruiz-Garbajosa, Rafael Cantón, Rafael Peña-Miller, Álvaro San Millán

**Author notes:** Correspondence: Álvaro San Millán, and Rafael Peña-Miller.

## Abstract

Plasmid persistence in bacterial populations is strongly influenced by the fitness effects associated with plasmid carriage. However, plasmid fitness effects in wild-type bacterial hosts remain largely unexplored. In this study, we determined the distribution of fitness effects (DFE) for the major antibiotic resistance plasmid pOXA-48 in wild-type, ecologically compatible enterobacterial isolates from the human gut microbiota. Our results show that although pOXA-48 produced an overall reduction in bacterial fitness, the DFE was dominated by quasi-neutral effects, and beneficial effects were observed in several isolates. Incorporating these data into a simple population dynamics model revealed a new set of conditions for plasmid stability in bacterial communities, with plasmid persistence increasing with bacterial diversity and becoming less dependent on conjugation. Moreover, genomic results showed a link between plasmid fitness effects and bacterial phylogeny, helping to explain pOXA-48 epidemiology. Our results provide a simple and general explanation for plasmid persistence in natural bacterial communities.

## Introduction

Plasmids are extra-chromosomal mobile genetic elements able to transfer between bacteria through conjugation^1^. Plasmids carry accessory genes that help their hosts to adapt to a myriad of environments and thus play a key role in bacterial ecology and evolution^2^. A key example of the importance of plasmids in bacterial evolution is their central role in the spread of antibiotic resistance mechanisms among clinical pathogens over recent decades^3,4^. Some of the most clinically relevant resistance genes, such those encoding carbapenemases (ß-lactamase enzymes able to degrade carbapenem antibiotics), are carried on conjugative plasmids that spread across high-risk bacterial clones^5,6^.

Despite the abundance of plasmids in bacterial populations and the potential advantages associated with their acquisition, these genetic elements generally produce physiological alterations in their bacterial hosts that lead to a reduction in fitness^7–9^. These fitness costs make it difficult to explain how plasmids are maintained in bacterial populations over the long-term in the absence of selection for plasmid-encoded traits, a puzzle known as “the plasmid-paradox”^10^. Different solutions to this paradox have been proposed. For example, compensatory evolution contributes to plasmid persistence by alleviating the costs associated with plasmid-carriage, and a high conjugation rate can promote the survival of plasmids as genetic parasites^11–18^.

Over the past decades, many studies have investigated the existence conditions for plasmids in bacterial populations^14,18–23^. However, understanding of plasmid population biology is held in check by limitations of the model systems used for its study. First, most experimental reports of fitness costs have studied arbitrary associations between plasmids and laboratory bacterial strains^7,24^. These examples do not necessarily replicate plasmid fitness effects in natural bacterial hosts, which remain largely unexplored. Second, studies tend to analyse the fitness effects of a single plasmid in a single bacterium. However, plasmid fitness effects can differ between bacteria^25–28^, and this variability may impact plasmid persistence in bacterial communities (for a relevant example see^29^). Third, most mathematical models of plasmid population biology study clonal or near-clonal populations. However, bacteria usually live in complex communities in which conjugative plasmids can spread between different bacterial hosts^30–32^. To fully understand plasmid persistence in natural bacterial populations, it will be necessary to address these limitations.

In this study, we provide the first description of the distribution of fitness effects (DFE) of a plasmid in wild-type bacterial hosts. We used the clinically relevant carbapenem-resistance conjugative plasmid pOXA-48 and 50 enterobacteria strains isolated from the gut microbiota of patients admitted to a large tertiary hospital in Madrid. Incorporation of the experimentally determined DFE into a population biology model provides new key insights into the existence conditions of plasmids in bacterial communities.

## Results

### Construction of a pOXA-48 transconjugant collection

We studied the DFE of the plasmid pOXA-48 in a collection of ecologically compatible bacterial hosts. pOXA-48 is an enterobacterial, broad-host-range, conjugative plasmid that is mainly associated with *K. pneumoniae* and *Escherichia coli*^33–35^. pOXA-48 encodes the carbapenemase OXA-48 and is distributed worldwide, making it one of the most clinically important carbapenemase-producing plasmids^6,34^. The gut microbiota of hospitalised patients is a frequent source of enterobacteria clones carrying pOXA-48^6^. In recent studies, we described the in-hospital epidemiology of pOXA-48 in a large collection of extended-spectrum ß-lactamase (ESBL)- and carbapenemase-producing enterobacteria isolated from more than 9,000 patients in our hospital over a period of two years (R-GNOSIS collection, see methods)^31,36–38^. pOXA-48-carrying enterobacteria were the most frequent carbapenemase-producing enterobacteria in the hospital, with 171 positive isolates, and they colonised 1.13% of the patients during the study period (105/9,275 patients). In this study we focused on plasmid pOXA-48_K8, which is a recently described pOXA-48-like plasmid isolated from a *K. pneumoniae* in our hospital^31^ (Figure 1a, for simplicity we will refer to pOXA-48_K8 and pOXA-48-like plasmids as pOXA-48 throughout the text).

**Figure 1.**
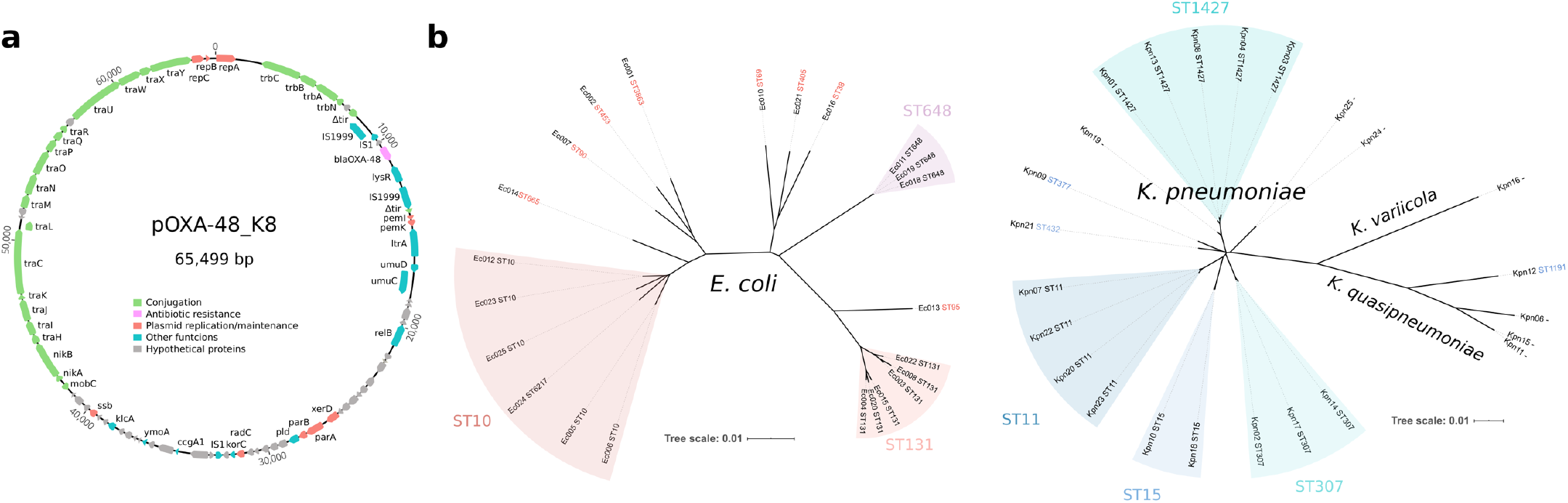
Experimental model system. Representation of pOXA-48 plasmid and the enterobacteria strains used in this study. (a) pOXA-48_K8 (accession number MT441554). Reading frames are shown as arrows, indicating the direction of transcription. Colours indicate gene function classification (see legend). The *bla*_OXA-48_ gene is shown in pink. (b) Unrooted phylogeny of whole-genome assemblies from *E. coli* clones (left) and *Klebsiella* spp. clones (right). Branch length gives the inter-assembly mash distance (a measure of k-mer similarity). The grouping of multi-locus sequence types (ST) is also indicated (*E. coli* ST6217 belongs to the ST10 group). Note that the sequencing results revealed that a subset of isolates initially identified as *K. pneumoniae* were in fact *Klebsiella quasipneumoniae* (n= 4) and *Klebsiella variicola* (n= 1).

To study the DFE of pOXA-48, we selected 50 isolates from the R-GNOSIS collection as bacterial hosts. Our criteria were to select (i) pOXA-48-free isolates, to avoid selecting clones in which compensatory evolution had already reduced plasmid-associated costs; (ii) isolates from the most frequent pOXA-48-carrying species, *K. pneumoniae* and *Escherichia coli;* and (iii) strains isolated from patients located in wards in which pOXA-48-carrying enterobacteria were commonly reported^31^. The underlying rationale was to select clones which were naïve to pOXA-48 but ecologically compatible with it *(i.e*. isolated from patients coinciding on wards with others who were colonised with pOXA-48-carrying clones). We selected 25 *K. pneumoniae* and 25 *E. coli* isolates that are representative of the R-GNOSIS study and cover the *K. pneumoniae* and *E. coli* phylogenetic diversity in the collection (see methods, Figure 1b and Supplementary Table 1). It is important to note that, because of the nature of the R-GNOSIS collection, the isolates used in this study produce ESBLs. However, ESBL-producing enterobacteria are widespread not only in hospitals but also in the community^39^, and most pOXA-48-carrying enterobacteria isolated in our hospital also produce ESBLs^36^.

pOXA-48 was introduced into the collection of recipient strains by conjugation (see Methods), and the presence of the plasmid was confirmed by PCR and antibiotic susceptibility testing (Supplementary Table 2). The presence of the entire pOXA-48 plasmid was confirmed by sequencing the complete genomes of the 50 transconjugant clones, which also revealed the genetic relatedness of the isolates (Figure 1b). In line with previous studies^31,40^, the sequencing results revealed that a subset of isolates initially identified as *K. pneumoniae* in fact belonged to the species *Klebsiella quasipneumoniae* (n= 4) and *Klebsiella variicola* (n= 1). These species are also pOXA-48 hosts in our hospital^31^ and so were maintained in the study (Figure 1b).

### Measuring pOXA-48 fitness effects

To measure pOXA-48 fitness effects, we performed growth curves and competition assays for all the plasmid-carrying and plasmid-free clones in the collection. We first performed growth curves in pure cultures to calculate maximum growth rate (μ_max_) and maximum optical density (OD_max_), which can be used to estimate the intrinsic population growth rate (*r*) and carrying capacity (*K*), respectively (Supplementary Figure 1). We also measured the area under the growth curve (AUC), which integrates information about *r* and *K*. To estimate plasmid-associated fitness effects, we compared these parameters between each plasmid-carrying and plasmid-free pair of isogenic isolates (Figure 2a). The results showed that, as expected, pOXA-48 produced an overall decrease in the parameters extracted from the growth curves. However, in a substantial subset of clones, plasmid acquisition was not associated with a reduction in these parameters (Figure 2a).

**Figure 2.**
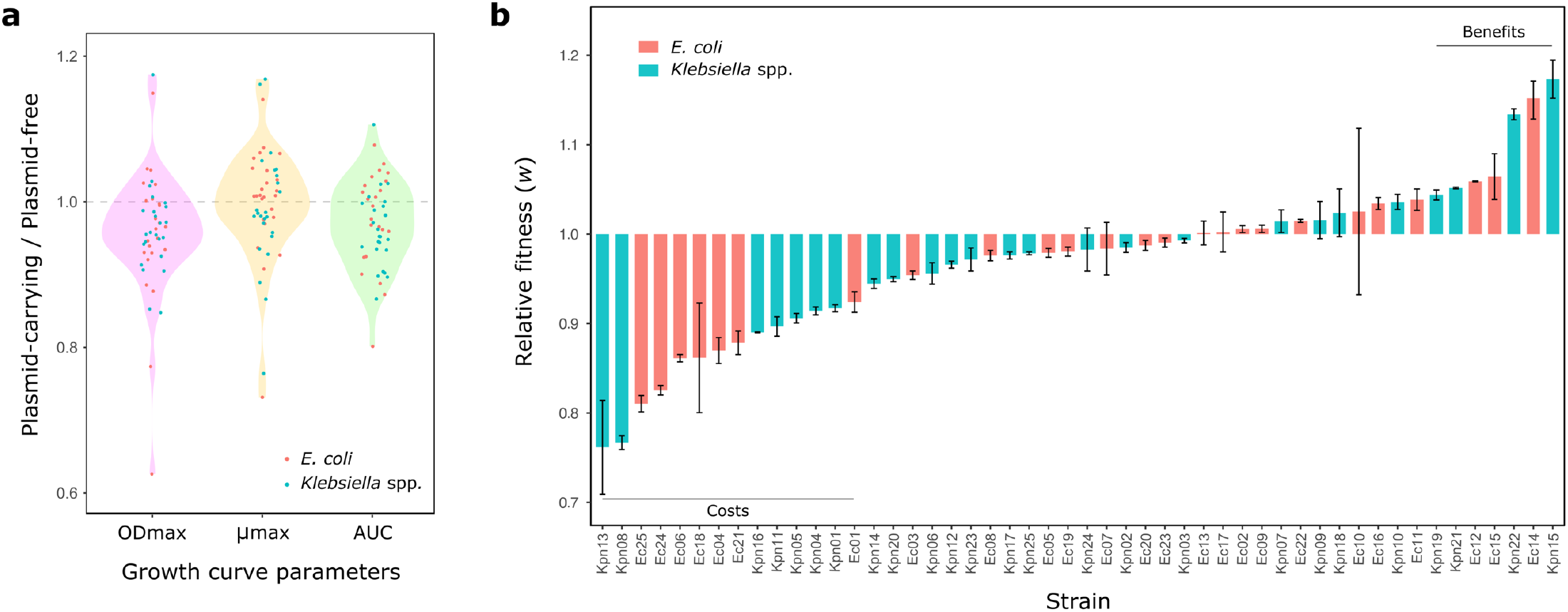
pOXA-48 fitness effects in a set of ecologically compatible wild-type enterobacteria. (a) Relative values of growth-curve parameters (plasmid-carrying/plasmid-free isogenic clones): maximum optical density (OD_max_, pink), maximum growth rate (μ_max_, yellow), and area under the curve (AUC, green). Dots represent each relative value (red, *E*. coli; blue, *Klebsiella* spp.). Values below 1 indicate a reduction in these parameters associated with plasmid-acquisition. Five biological replicates were performed for each growth curve. (b) Relative fitness (*w*) of plasmid-carrying clones compared with plasmid-free clones obtained by competition assays (red, *E. coli;* blue, *Klebsiella* spp.). Values below 1 indicate a reduction in *w* due to plasmid acquisition; values above 1 indicate an increase in *w*. Bars represent the mean of five independent experiments, and error bars represent the standard error of the mean. Two horizontal lines indicate those clones showing significant costs or benefits associated with carrying pOXA-48 plasmid.

Competition assays allow measurement of the relative fitness *(w)* of two bacteria competing for resources in the same culture^41^. Competition between otherwise isogenic plasmid-carrying and plasmid-free clones thus provides a quantitative assessment of the fitness costs associated with plasmid carriage. For the competition assays, we used flow cytometry; strains were labelled using an in-house developed small, non-conjugative plasmid vector, called pBGC, that encodes an inducible green fluorescent protein (GFP) (Supplementary Figure 2). pBGC was introduced into the wild-type isolate collection by electroporation, and all pOXA-48-carrying and pOXA-48-free clones were competed against their pBGC-carrying parental strain. We were unable to introduce pBGC into eight of the isolates; in those cases, for the competitor, we used *E. coli* strain J53 carrying the pBGC vector (see Methods for details). Data from the competition assays were used to calculate the competitive fitness of pOXA-48-carrying clones relative to their plasmid-free counterparts (Figure 2b). There were no significant differences between the fitness effects of pOXA-48 in *Klebsiella* spp. and *E. coli* isolates (ANOVA effect of Species x Plasmid interaction; F=0.088, df=1, P=0.767).

To validate our results, we compared the values obtained from growth curves and competition assays. This analysis revealed a significant correlation between relative fitness values and the parameters extracted from the growth curves (Supplementary Figure 3).

### The distribution of pOXA-48 fitness effects

Results from the competition assays showed that the overall effect of pOXA-48 was a small but significant reduction in relative fitness (mean *w*= 0.971, ANOVA effect of plasmid; F=70.04, df=1, P=1.02×10^−15^). However, plasmid fitness effects varied greatly between the isolates in the collection, producing a normal distribution ranging from a >20% reduction to almost a 20% increase in relative fitness (Figure 2b and 3a; Shapiro-Wilk normality test, P= 0.14). Indeed, plasmid acquisition was associated with a significant fitness decrease in only 14 strains, and 7 isolates showed a significant increase in fitness (Bonferroni corrected two sample t-test, *P*< 0.05). These results revealed a highly dynamic scenario in which a plasmid produces a wide distribution of fitness effects in different bacterial hosts, ranging from costs to benefits.

**Figure 3.**
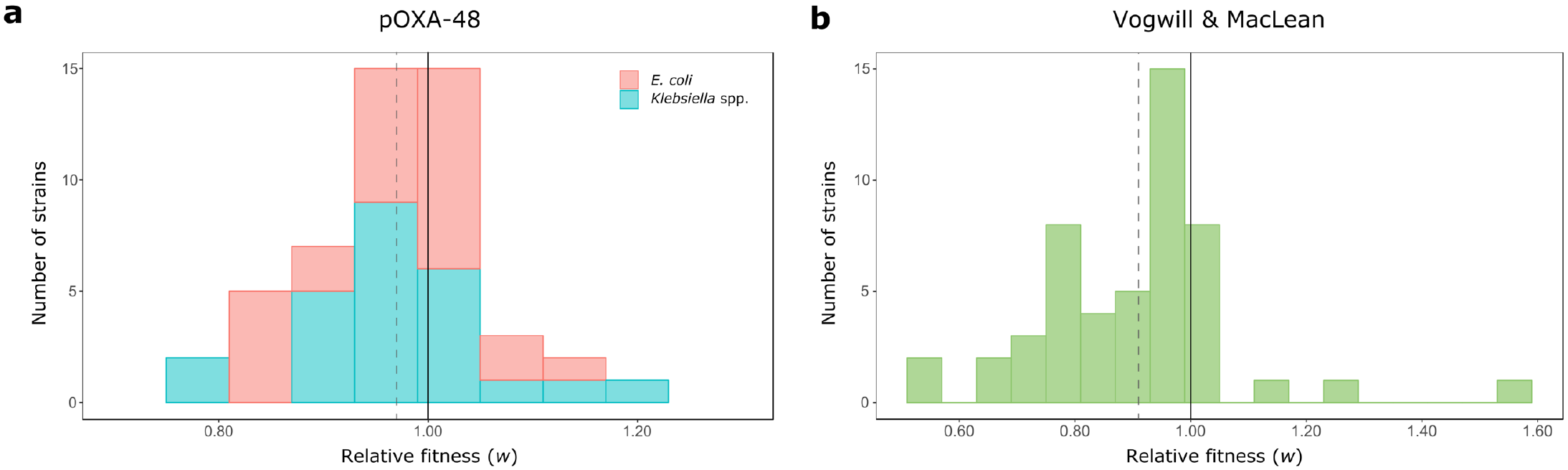
Distribution of plasmid fitness effects. Comparison between the DFE obtained in this study and the DFE from previous studies. (a) DFE for pOXA-48 in the ecologically compatible collection of enterobacteria isolates. Bars indicate the number of *E. coli* (red) and *Klebsiella* spp. (blue) strains in each relative fitness category. The grey dotted line indicates the mean relative fitness of the population. Note that relative fitness values are normally distributed (*w*= 0.971, var= 0.0072). (b) DFE for plasmids in bacterial hosts obtained in a previous meta-analysis^24^. Most of the included studies were based on arbitrary associations between plasmids and laboratory strains. Bars indicate the number of plasmid-bacterium associations in each relative fitness category. The grey dotted line indicates the mean relative fitness across studies. Relative fitness values are not normally-distributed (*w*= 0.91, var= 0.029; Shapiro-Wilk normality test, P= 0.0006).

To place our results in context with previous reports, we compared the DFE for pOXA-48 with the results from a recent meta-analysis of plasmid fitness effects by Vogwill and MacLean^24^ (Figure 3). These authors recovered data for 50 plasmid-bacterium pairs from 16 studies. The DFE constructed from those reports showed a higher mean plasmid cost (mean *w*= 0.91) and differed significantly from the DFE we report here for pOXA-48 in wild-type enterobacteria (Wilcoxon signed rank test, V= 922, P= 0.006). The discrepancy between these distributions may, at least in part, reflect the different nature of plasmid-bacterium associations considered in the different studies. Although the plasmids studied in earlier reports were isolated from natural sources, they were introduced into laboratory bacterial strains, and the detected fitness effects may not be fully representative of wild-type plasmid-bacterium associations. Our study, on the other hand, analysed the fitness effects of pOXA-48 in ecologically compatible bacterial hosts. Taken together, the data suggest that the distribution of plasmid fitness effects is likely influenced by the ecological compatibility between plasmids and their bacterial hosts.

### pOXA-48 fitness effects across bacterial phylogeny

A key limit to the prediction of plasmid-mediated evolution is the inability to anticipate plasmid fitness effects in new bacterial hosts. This is particularly relevant to the evolution of antibiotic resistance because some of the most concerning multi-resistant clinical pathogens arise from very specific associations between resistance plasmids and high-risk bacterial clones^4,6,42^. Interestingly, a recent study in an important pathogenic *E. coli* lineage (ST131) showed that the acquisition and maintenance of resistance plasmids is associated with specific genetic signatures^43^. Pursuing this idea, we analysed the DFE for pOXA-48 across the whole-genome phylogeny of our isolates, with the aim of determining if genetic content could help to predict plasmid fitness effects (Figure 4). We calculated the genetic relatedness of *Klebsiella* spp. and *E. coli* isolates by reconstructing their core genome phylogeny (Figure 4a). Plasmid fitness effects can also be strongly influenced by the accessory genome. For example, the presence of further mobile genetic elements can deeply impact the costs of plasmids^44,45^. Therefore, we also constructed trees from the distance matrix of the accessory gene network^46^, which includes plasmid content (Figure 4b).

**Figure 4.**
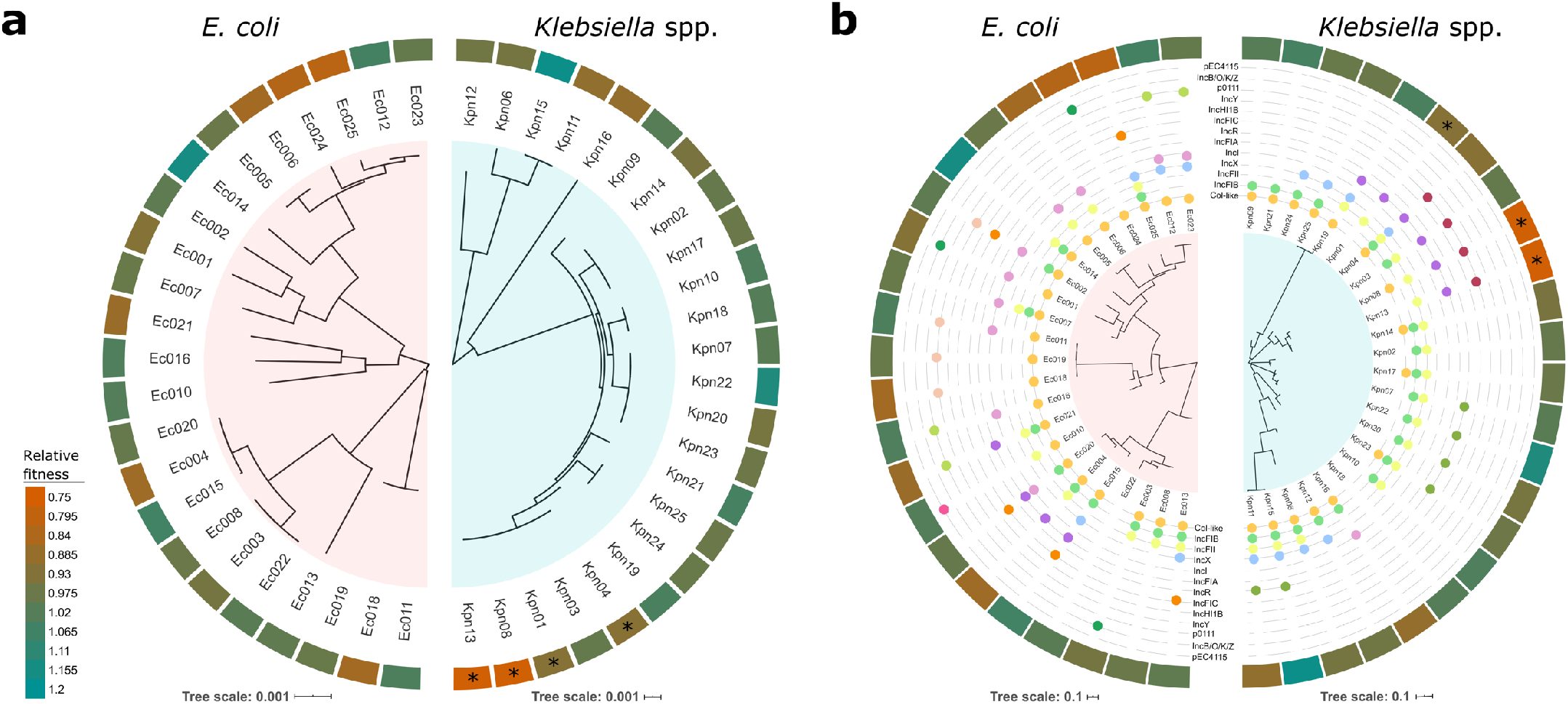
Fitness effects of pOXA-48 across bacterial genome content. An association was found between pOXA-48 fitness effects and bacterial host genomic content for four *K. pneumoniae* ST1427 isolates. (a) Core genome relationships among *E. coli* (left) and *Klebsiella* spp. (right). Tree construction is based on polymorphisms in the core genome. The outer circle indicates the relative fitness of pOXA-48-carrying bacterial hosts (see legend for colour code; red indicates fitness costs and green indicates fitness benefits associated with pOXA-48 carriage). Asterisks denote clones with a phylogenetic signal associated with plasmid fitness effects (LIPA, P< 0.05). (b) Accessory genome relationships among *E. coli* (left) and *Klebsiella* spp. (right). Tree construction is based on the distance matrix of the accessory gene network of each group. The outermost circle indicates relative fitness as in (a). The intermediate circles indicate presence/absence of plasmids belonging to the different plasmid families named in the figure. Asterisks denote clones with a significant phylogenetic signal associating accessory genome composition with pOXA-48 fitness effects (LIPA, P< 0.05).

For each group of isolates, we scanned the fitness effects of pOXA-48 across the core and accessory genome using the local indicator of phylogenetic association index^47,48^ (LIPA, see Supplementary Figure 4, Supplementary Table 3, and methods for the complete analysis). For the *E. coli* isolates, the results showed no association of pOXA-48 fitness effects with the core or accessory phylogenies (LIPA, P> 0.1). In contrast, for *Klebsiella* spp., LIPA indices revealed a significant phylogenetic signal in four clones in which pOXA-48 produced a high fitness cost, all of them belonging to ST1427 (Kpn01, Kpn04, Kpn08, and Kpn13, accounting for 4 of the 5 ST1427 clones analysed in this study; LIPA, P< 0.05). Three of these ST1427 clones also produced a significant signal in the analysis of fitness effects across the accessory genome (Kpn01, Kpn08, and Kpn13; LIPA, P< 0.05). The results thus reveal that pOXA-48 tended to produce a high cost in *K. pneumoniae* clones belonging to ST1427. Interestingly, although *K. pneumoniae* ST1427 is relatively common in our hospital (4.8% of ESBL-producing *K. pneumoniae*^38^), none of the 103 pOXA-48-carrying *K. pneumoniae* isolates recovered in the R-GNOSIS collection belong to this ST^31^ (Fisher’s exact test for count data, 8/166 vs 0/103, P= 0.025). These results suggest that the high cost associated with plasmid acquisition in this clade may limit in-hospital spread of pOXA-48-carrying *K. pneumoniae* ST1427. Conversely, pOXA-48 is commonly associated with *K. pneumoniae* ST11 in our hospital^31,36^, and in the four ST11 clones tested in this study, pOXA-48 produced neutral (Kpn07, Kpn20, Kpn23) or even beneficial fitness effects (Kpn22, Figure 4A) (pOXA-48 fitness effects in ST1427 [n=5] vs. in ST11 [n=4], Welch’s unequal variances two-tailed t-test, t= - 2.39, df= 7, P= 0.048).

### Modelling the role of DFE in plasmid stability

In general, mathematical models of plasmid population biology consider a clonal population in which the plasmid produces a constant reduction in growth rate^14,18–23^. These models usually include the rate of plasmid loss through segregation^49,50^ and the rate of horizontal plasmid transfer by conjugation^19,20,51^, and some of them also incorporate a rate of compensatory mutations that alleviate plasmid fitness costs over time^14,23^. Our results show that plasmids produce a wide DFE in naturally compatible bacterial hosts, and this distribution could strongly influence plasmid stability in polyclonal bacterial communities. To assess the effect of the DFE on plasmid stability in bacterial communities, we developed a simple mathematical model based on Stewart and Levin’s pioneering work on plasmid existence conditions^19^.

The model describes the population dynamics of multiple subpopulations competing for a single exhaustible resource in well-mixed environmental conditions, assuming that transition between plasmid-bearing and plasmid-free cells is driven by segregation events. The growth rate of each subpopulation is determined by a substrate-dependent Monod term that depends on the extracellular resource concentration, and therefore each strain can be described by two structurally identifiable parameters^49^: the resource conversion rate (ρ) and the specific affinity for the resource (V_max_/*K*_m_). These parameters were estimated from the optical densities of each strain growing in monoculture (with and without plasmids) using a Markov chain Monte Carlo (MCMC) method with a Metropolis-Hastings sampler (See Methods, Figure 5a and Supplementary Figure 5).

**Figure 5.**
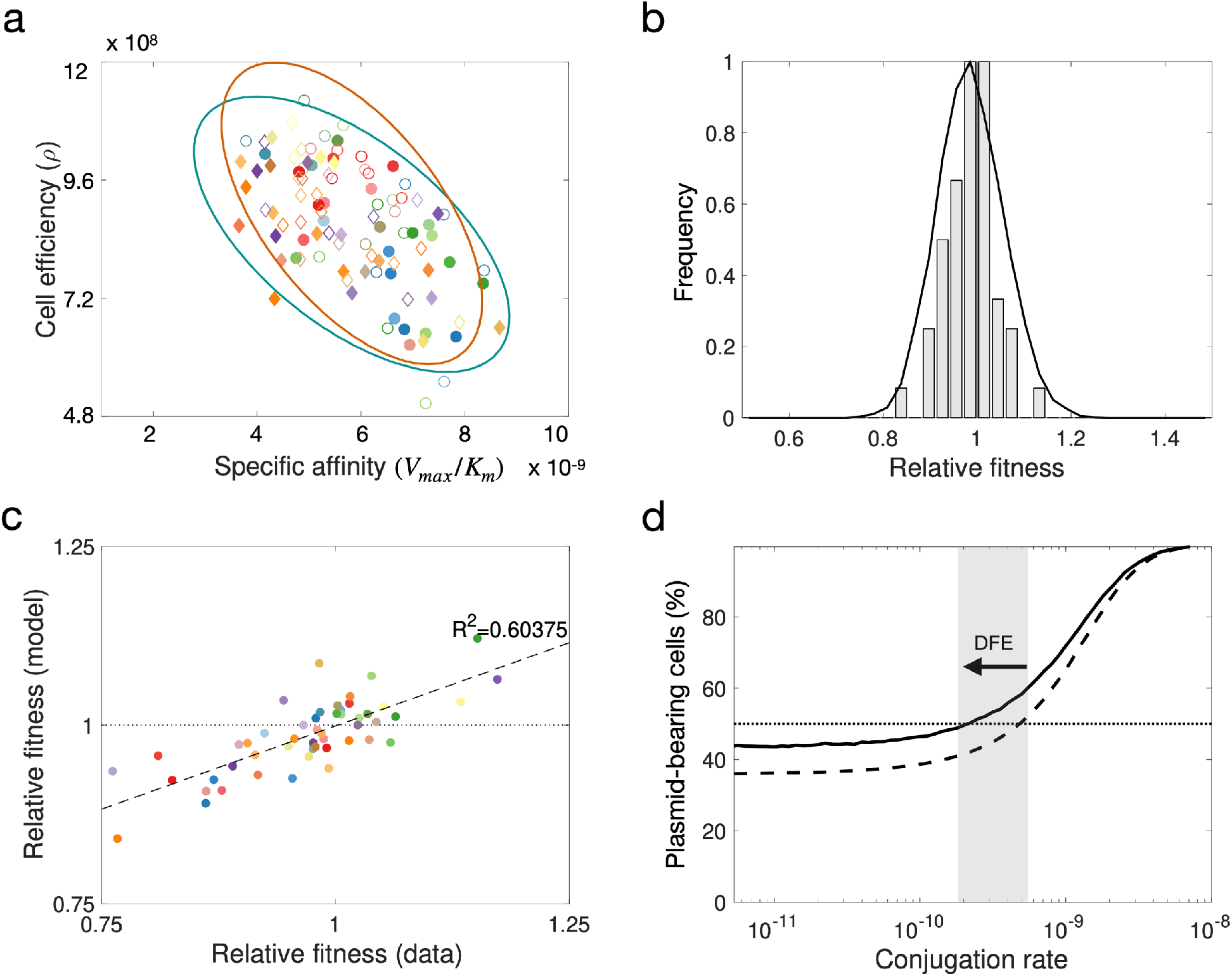
Modelling the DFE for pOXA-48. (a) Distribution of parameter values obtained using Bayesian inference to estimate growth kinetic parameters from OD measurements obtained for each strain in isolation. Diamonds represent *Klebsiella* spp. strains and circles *E. coli* clones; filled symbols denote plasmid-bearing strains and empty symbols plasmid-free cells. The ellipses represent standard deviations of best-fit Normal distributions (green for plasmid-bearing strains and orange for plasmid-free cells). (b) Bars represent a DFE obtained from *in silico* competition experiments with parameter values determined from experimental growth curves. The solid curve represents the computationally estimated DFE obtained by randomly sampling wild-type and transconjugant parameter distributions obtained using the MCMC algorithm and numerically solving the model to evaluate the relative fitness associated with plasmid carriage. (c) Comparison of relative fitness values obtained experimentally and using the population dynamics model (R^2^= 0.603). (d) Fraction of plasmid-bearing cells as a function of the rate of horizontal transfer for random plasmid-host associations sampled from the MCMC parameter distribution. The dotted line illustrates the mean of 10^4^ pair-wise competition experiments under the assumption plasmid-bearing is associated with a constant reduction in fitness in different clones (*w*= 0.985, var= 0), while the solid line is obtained by considering a wide DFE (*w*= 0.985, var= 0.0070). The arrow denotes the difference in the conjugation threshold that positively selects for plasmids in the population, supporting the tenet that the DFE maintains plasmids in the population at lower conjugation rates.

By solving the system of differential equations (described in Methods), we were able to evaluate the final frequency of plasmid-bearing cells in an experiment of duration *T* time units and quantify the fitness effect of the plasmid on the strain. Figure 5b shows the DFE obtained after performing *in silico* pair-wise competition experiments between plasmid-bearing and plasmid-free subpopulations (with parameter values shown in Supplementary Table 5, Supplementary Figure 6), resulting in a theoretical DFE (*w*= 0.985, var= 0.0070) that is consistent with the experimentally measured DFE presented in Figure 3 (*w*= 0.971, var= 0.0072). Moreover, comparison of model predictions with relative fitness values obtained by flow cytometry are consistent (R^2^= 0.603; Figure 5c), showing that the population dynamics model can accurately predict the outcome of a competition experiment from the individual growth dynamics.

Previous studies showed that the probability of plasmid fixation is correlated with the rate of horizontal transmission^19,20,49^. As previous models, we consider horizontal transmission of plasmids as a function of the densities of donor and recipient cells, with conjugation events occurring at a constant rate. Competition experiments for a range conjugation rates are illustrated in Figure 5d; while at low horizontal transmission rates plasmid-free cells outcompete plasmid-bearing cells, at higher conjugative rates, plasmid-bearing cells increase in frequency.

### Community complexity promotes plasmid persistence

To explore how plasmid stability is affected by increasing community complexity and rates of horizontal transmission, we randomly sampled *N*= 10^4^ plasmid-free cells from the distribution of growth parameters estimated using the MCMC algorithm. These random communities were used to study the population dynamics of plasmids transmitting vertically and horizontally in multi-strain communities. The fitness cost (or benefit) of bearing plasmids was modelled as a random variable that modifies the wild-type (plasmid-free) growth rate by a factor σ, such that if σ= 0, the DFE has zero variance (Figure 6a), but if σ> 0, the resulting DFE is a symmetrical heavy-tailed distribution with a right-hand tail expanding towards positive fitness effects (Figure 6b), indicating the existence of plasmid-host associations in which plasmid carriage produces a fitness benefit.

**Figure 6.**
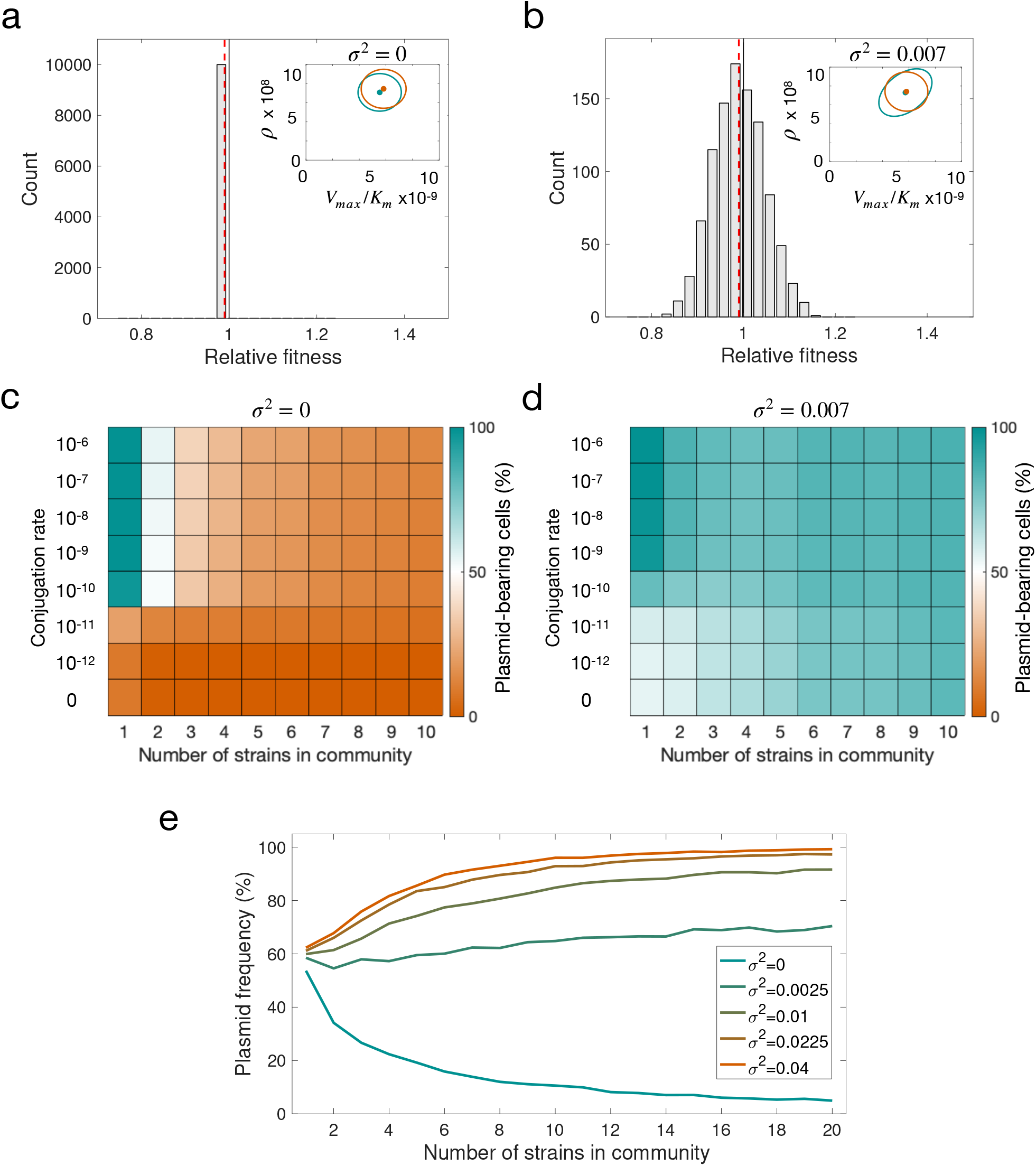
Modelling plasmid persistence in polymicrobial communities, assuming fixed (a,c) or variable (b,d) plasmid fitness effects. (a-b) Relative fitness histogram obtained by randomly sampling 10^4^ parameter values from the parameter distribution shown in the inset plot (points illustrate the expected values of each distribution and ellipses their standard deviation; green, plasmid-bearing bacteria; orange, plasmid-free bacteria). The green ellipse in b is larger as a consequence of considering that the cost of plasmid-bearing is normally-distributed with variance 0.007. As a result, the DFE also has higher variance, with a considerable fraction of plasmid-host associations producing a benefit to the host. Dotted red lines indicate mean relative fitness of plasmid carrying cells. (c-d) Colour gradient represents the percentage of cells carrying plasmids at the end of 5,000 stochastic simulations; orange indicates a population without plasmids and green a community composed of plasmid-carrying cells. If plasmid-bearing is associated with a fixed fitness cost for all members of the community, plasmid maintenance requires a high conjugation rate. The increased proportion of plasmid-bearing cells in d indicates that a DFE with high variance reduces the critical conjugation rate needed to maintain plasmids in the population, enabling plasmids to persist at low conjugation rates. (e) Mean fraction of plasmid-bearing cells as a function of the number of strains in the community with a conjugation rate γ= 1.5 × 10^−11^. If the plasmid always produces a reduction in host fitness (mean *w* <1 and low variance), plasmid frequency decreases as the number of strains in the community increases (green line). In contrast, for higher variance at the same mean *w*, the fraction of plasmid-bearing cells increases with community complexity (orange line).

To assess how DFE influences plasmid persistence in polymicrobial communities, we extended the model to consider populations composed of subsets of 1, 2, 3, 4, …, *M* ≤ *N* cell types sampled randomly from the wild-type parameter distribution (see Methods and Supplementary Figure 7 and 8). This enabled us to estimate the relative frequency of plasmid-bearing cells at the end of a long-term experiment and evaluate the stability of the plasmid in multi-strain communities with different population structures. Initial bacterial densities were determined by first running the system forward (with all strains initially present at equal densities and carrying pOXA-48) for *T*= 24 time units, and then clearing all plasmid-free cells from the population. This assumption is akin to patients receiving an antibiotic therapy that clears all plasmid-free cells from the bacterial community. The results obtained after 5,000 computer simulations over a range of conjugation rates and numbers of cell types in the community are shown in Figure 6. The simulations either assumed an identical fitness cost for all strains (*w*= 0.985, var= 0; Figure 6a,c) or allowed plasmid fitness effects to vary according to the experimentally determined DFE (w= 0.985, var= 0.0070; Figure 6b,d).

Although the mean fitness cost was the same in both conditions, the results of the computational experiments suggest that allowing fitness effects to vary between members of the population markedly increases the chances of plasmid persistence, especially at low conjugation rates. More importantly, in the numerical simulations, plasmid frequency decreased as a function of the number strains in the community when plasmid acquisition was associated with a constant fitness cost, but increased with community complexity for DFEs with larger variance (Figure 6c,d,e). The explanation for this effect is that if plasmid fitness cost is identical for all community members, diversity simply means extra competition for plasmid-carrying cells, and plasmid persistence becomes more dependent on a high conjugation rate. In contrast, if the fitness effects vary, a larger number of available bacterial hosts in the population increases the probability of the plasmid arriving to a host in which it produces a neutral or beneficial fitness effect. This is an important result because it implies that increasing bacterial community complexity could increase the probability of plasmid persistence in natural environments. Given that most natural microbiota are complex and plasmids can usually conjugate and replicate in different clones, this may explain the high prevalence of plasmids in nature. Our results also indicate that the threshold conjugation rate for plasmid persistence may be lower than previously thought. In fact, once plasmids are present in multiple members of a community, they may be able to persist even in the absence of conjugation (Figure 6d).

## Discussion

The DFE for new mutations is a central concept in genetics and evolutionary biology, with implications ranging from population adaptation rates to complex human diseases^52^. The fitness effects of new spontaneous mutations in bacteria follow a heavy-tailed distribution dominated by quasi-neutral mutations with infrequent strongly deleterious mutations^53,54^. Horizontally acquired genes also produce a distribution of fitness effects in new bacterial hosts^55,56^. However, horizontal gene transfer in bacteria is frequently mediated by entire mobile genetic elements, such as plasmids, that carry multiple genes. Numerous studies have measured the fitness effects of individual plasmids in a bacterial host^24^, but the DFE of a plasmid in multiple, ecologically compatible bacterial hosts had not been reported before. Here, we determined the DFE of a carbapenem resistance plasmid in wild-type enterobacteria recovered from the human gut microbiota. Unsurprisingly, the DFE of pOXA-48 differed from the DFE of spontaneous mutations. As with spontaneous mutations, the pOXA-48 DFE was also dominated by quasi-neutral effects and was slightly shifted towards fitness costs; however, instead of a single heavy tail of deleterious effects, it had a symmetrical shape, with tails expanding both towards negative and positive fitness effects (Figure 3a).

Two key implications of the experimentally determined DFE in this study are that, according to our simple mathematical model, the probability of plasmid persistence becomes less dependent on a high conjugation rate and increases with the number of bacterial strains in the population. The complex and multi-clonal nature of most natural bacterial communities attests the likely relevance of our findings to the extremely high prevalence of plasmids in bacterial populations^57^. The human gut microbiota, for example, includes a great variety of bacteria from hundreds of species^58^, including several strains from the *Enterobacterales* order alone^59^. Our experimental system is in fact inspired by the dynamics of pOXA-48 in the gut microbiota of hospitalised patients. In a recent study, we observed that once patients are colonised by a pOXA-48-carrying clone, the plasmid spreads through conjugation to other resident enterobacteria present in the gut microbiota^31^. Crucially, pOXA-48 usually persists in the gut of patients throughout the hospital stay and can be detected in subsequent hospital admissions months or years later, and not necessarily in the original colonizing strain^31^. Our results indicate that the pOXA-48 DFE could explain the long-term persistence of this and other plasmids in the human gut microbiota.

Another interesting result of this study is that pOXA-48 produced a particularly elevated cost in *K. pneumoniae* isolates belonging to ST1427 (Figure 4A). ST1427 is under-represented among the pOXA-48-carrying *K. pneumoniae* isolates in our hospital, which are dominated by ST11^31,36^. Remarkably, in the four *K. pneumoniae* ST11 clones tested in this study, pOXA-48 produced neutral (Kpn07, Kpn20, Kpn23) or even beneficial fitness effects (Kpn22, Figure 4A). Therefore, despite the small number of *K. pneumoniae* clones analysed, our results suggest that phylogeny might influence fitness compatibility between plasmids and bacteria at the clonal level, dictating the epidemiology of plasmid-bacterium associations in clinical settings. Further analysis of a larger sample of *K. pneumoniae* isolates from the different STs will be needed to elucidate the genetic basis underlying these specific interactions between bacterial phylogeny and pOXA-48 fitness effects.

The main experimental limitation of our study is that plasmid fitness effects were determined *in vitro*, using planktonic cultures in LB medium. This is the standard practise in the field, and previous studies have shown that plasmid fitness effects measured in laboratory conditions correlate with those measured in animal models^24^; however, our results may not be fully representative of pOXA-48 fitness effects in the human gut. Future studies will need to explore more complex *in vitro* systems^60^, as well as *in vivo* animal models^61^. Another important limitation of our study is that we modelled bacterial communities with a simple resource competition model that does not consider spatial structure^62^, complex ecological interactions between community members^63^, plasmid-host co-evolution^64^, or differential rates of horizontal transmission^28^. Although more complex models^65^ will be needed to integrate bacterial community complexity and plasmid fitness effects, consideration of diverse polymicrobial populations with complex spatiotemporal interactions would likely only increase DFE variance, therefore promoting plasmid stability.

## Methods

### Strains, pOXA-48 plasmid, and culture conditions

We selected 50 representative ESBL-producing clones form the R-GNOSIS collection (Supplementary Table 1). This collection was constructed in our hospital as part of an active surveillance-screening program for detecting patients colonised by ESBL/carbapenemase-producing enterobacteria, from March 4th, 2014, to March 31^st^, 2016 (R-GNOSIS-FP7-HEALTH-F3-2011-282512, www.r-gnosis.eu/, approved by the Ramón y Cajal University Hospital Ethics Committee, Reference 251/13)^36,38^. The screening included a total of 28,089 samples from 9,275 patients admitted at 4 different wards (gastroenterology, neurosurgery, pneumology and urology) in the Ramon y Cajal University Hospital (Madrid, Spain). The characterisation of samples was performed during the R-GNOSIS study period^36,66^; rectal swabs were plated on Chromo ID-ESBL and Chrom-CARB/OXA-48 selective agar media (BioMérieux, France) and bacterial colonies able to grow on these media were identified by MALDI-TOF MS (Bruker Daltonics, Germany) and further characterized by pulsed-field gel electrophoresis (PFGE). For the present study, we selected 25 *E. coli* and 25 *K. pneumoniae* ESBL-producing isolates from the R-GNOSIS collection. The strains were representative of *E. coli* and *K. pneumoniae* diversity in the R-GNOSIS collection (randomly chosen form the most common pulsed-field gel electrophoresis profiles^38^), they did not carry any carbapenemase gene and they were recovered from patients not colonised by other pOXA-48-carrying clones. To construct the transconjugants, we used the most common pOXA-48 plasmid variant from the R-GNOSIS collection in our hospital, according to plasmid genetic sequence (pOXA-48_K8, accession number MT441554)^30^. Bacterial strains were cultured in lysogeny broth (LB) at 37°C in 96-well plates with continuous shaking (250 rpm) and on LB agar plates at 37°C.

### Construction of transconjugants collection

We performed an initial conjugation round to introduce pOXA-48_K8 plasmid from wild type *E. coli* C609 strain^31^, into *E. coli* β3914^67^, a diaminopimelic acid (DAP) auxotrophic laboratory mutant of *E. coli* K-12 (kanamycin, erythromycin and tetracycline resistant, Supplementary Table 1), which was used as the common counter-selectable donor. The pOXA-48-carrying wild type *E. coli* C609 and *E. coli* β3914 were streaked from freezer stocks onto solid LB agar medium with ertapenem 0.5 μg/ml and DAP 0.3 mM, respectively, and incubated overnight at 37°C. Donor and recipient colonies were independently inoculated in 2 ml of LB in 15-ml culture tubes and incubated overnight. After growth, donor and recipient cultures were collected by centrifugation (15 min, 1,500 g) and cells were re-suspended in each tube with 300 μl of sterile NaCl 0.9%. Then, the suspensions were mixed in a 1:1 proportion, spotted onto solid LB medium with DAP 0.3 mM and incubated at 37°C overnight. Transconjugants were selected by streaking the conjugation mix on LB with ertapenem (0.5 μg/ml), DAP 0.3 mM, tetracycline (15 μg/ml), and kanamycin (30 μg/ml). The presence of pOXA-48 was checked by PCR, using primers for *bla*_OXA-48_ gene and for the replication initiation protein gene *repC* (Supplementary Table 4).

We used the counter-selectable *E. coli* β3914/pOXA-48_K8 donor to conjugate plasmid pOXA-48 in the 50 wild type strains. We used the same protocol described above, but the final conjugation mix was plated on LB with no DAP (to counter-select the donor) and with amoxicillin-clavulanic acid (to select for transconjugants). The optimal concentration of amoxicillin-clavulanic acid was experimentally determined for each isolate in the collection and ranged from 64 μg/ml to 384 μg/ml. The presence of pOXA-48 in the transconjugants was checked by PCR, as described above, and by antibiotic susceptibility testing and whole genome sequencing (see below). To test the stability of plasmid pOXA-48 in the transconjugants we propagated cultures in LB with no antibiotic selection (two consecutive days, 1:10,000 dilution) and plated cultures on LB agar. After ON incubation at 37°C, 100 independent colonies of each transconjugant were replicated both on LB agar and LB agar with amoxicillin-clavulanic acid to identify pOXA-48-carrying colonies (including negative controls of plasmid-free wild type clones). Results showed that the plasmid was overall stable in the transconjugants; 100% stable in 43 isolates, and ≥ 90% stable in the 7 remaining isolates.

### Antibiotic susceptibility testing

Antibiotic susceptibility profiles were determined for every wild-type and transconjugant strain by the disc diffusion method following the EUCAST guidelines (www.eucast.org) (Supplementary Table 2). We used the following antimicrobials agents: imipenem (10 μg), ertapenem (10 μg), amoxicillin-clavulanic acid (20/10 μg), rifampicin (30 μg), streptomycin (300 μg), chloramphenicol (30 μg) and amikacin (30 μg) (Bio-Rad, CA, USA). pOXA-48-carrying and pOXA-48-free strains were pre-cultured in Müller-Hinton (MH) broth at 37 °C in 15 ml test tubes with continuous shaking (250 rpm), and disc diffusion antibiograms were performed on MH agar plates (BBL, Becton Dickinson, MD, USA).

### Growth curves

Pre-cultures of plasmid-free and plasmid-carrying strains (5 replicates of each) were prepared by inoculating single independent colonies into LB broth and overnight incubation at 37 °C with continuous shaking (250 rpm). Overnight cultures were diluted 1:1,000 into fresh LB in 96-well plates, which were incubated during 22 h at 37 °C with shaking (250 rpm) in a plate reader (Synergy HTX Multi-Mode Reader, BioTek Instruments, Inc, VT, USA). Optical densities (OD) were measured every 15 minutes during the incubation. The maximum growth rate (μ_max_), maximum optical density (OD_max_), and area under the growth curve (AUC) were determined using Gen5™ Microplate Reader and Imager Software and the *growthrates* package in R. We calculated the relative OD_max_, μ_max_, and AUC, by dividing the average value of each parameter for the pOXA-48-carrying isolate between that of the pOXA-48-free isolate using the follow formula:

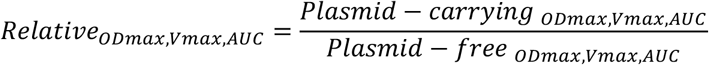

### Construction of pBGC, a GFP-expressing non-mobilizable plasmid

To fluorescently label the wild type isolates for competition assays using flow cytometry, we constructed the pBGC plasmid, a non-mobilizable version of the *gfp*-carrying small plasmid pBGT^68^ (Supplementary Figure 2, accession number MT702881). The pBGT backbone was amplified, except for the region including the oriT and *bla*TEM1 gene, using the pBGC Fw/Rv primers. The *gfp* terminator region was independently amplified using the GFP-Term Fw/Rv primers (Supplementary Table 4). PCR amplifications were made with Phusion Hot Start II DNA Polymerase at 2 U/μL (ThermoFisher Scientific, MA, USA), and PCR products were digested with DpnI to eliminate plasmid template before setting up the assembly reaction (New England BioLabs, MA, USA). Finally, pBGC was constructed by joining the amplified pBGT backbone and the *gfp* terminator region using the Gibson Assembly Cloning Kit (New England BioLabs, MA, USA). Resulting reaction was transformed by heat shock into NEB 5-alpha Competent *E. coli* (New England BioLabs, MA, USA), following manufacturer’s instructions. Transformation product was plated on LB agar with arabinose 0.1% and chloramphenicol 30 μg/ml, and incubated overnight at 37 °C. Plasmid-bearing colonies were selected by green fluorescence. The *gfp* gene in pBGC is under the control of the P_*BAD*_ promoter, so GFP production is generally repressed and induced by the presence of arabinose. pBGC was completely sequenced using primers described in Supplementary Table 4. We confirmed that neither pOXA-48, nor helper plasmid pTA-Mob^69^, could mobilized pBGC by conjugation using the conjugation protocol described above, confirming that pBGC plasmid is not mobilizable. Finally, pBGC plasmid was introduced into our isolate collection by electroporation (Gene Pulser Xcell Electroporator, BioRad, CA, USA). Of note, we were not able to obtain pBGC-carrying transformants in eight of the isolates due to a pre-existing high chloramphenicol resistance phenotype.

### Competition assays using flow cytometer

We performed competition assays^41^, using flow cytometry, to obtain the relative fitness of pOXA-48-carrying isolates compared to their pOXA-48-free parental counterparts. We used the collection of pBGC transformed wild type isolates as competitors against their isogenic pOXA-48-carrying and pOXA-48-free isolates. Specifically, two sets of competitions were performed for each isolate: pOXA-48-free vs. pBGC-carrying, and pOXA-48-carrying vs. pBGC-carrying. Five biological replicates of each competition were performed. Pre-cultures were incubated overnight in LB in 96-well plates at 225 rpm an 37°C, then mixed 1:1 and diluted 10,000-fold in 200 μl of fresh LB in in 96-well plates. Mixtures were competed for 24 h in LB at 37°C and 250 rpm (the low initial cell density and the strong shaking hinders pOXA-48 conjugation, see control experiment below). To determine the initial proportions, initial 1:1 mixes were diluted 2,000-fold in 200 μl of NaCl 0.9 % with L-arabinose 0.1 %, and incubated at 37 °C at 250 rpm during 1.5 h to induce *gfp* expression. The measurements were performed via flow cytometry using a CytoFLEX Platform (Beckman Coulter Life Sciences, IN, US) with the following parameters: 50 μl min^−1^ flow rate, 22 μm core size, and 10,000 events recorded per sample (Supplementary Figure 9). After 24 hours of incubation, final proportions were determined as described above, after 2,000-fold dilution of the cultures. The fitness of each strain relative to its pBGC-carrying parental isolate was determined using the formula:

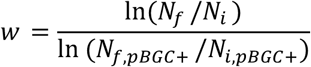

where *w* is the relative fitness of the pOXA-48-carrying (*w*_pOXA-48+_) or pOXA-48-free (*w*_pOXA-48−_) isolates compared to the pBGC-bearing parental clone, *N_i_* and *N_f_* are the number of cells of the pBGC-free clone at the beginning and end of the competition, and *N_i, pBGC_* and *N_f, pBGC_* are the number of cells of the pBGC-carrying clone at the beginning and end of the competition, respectively. The fitness of the pOXA-48-carrying isolates relative to the pOXA-48-free parental isolates were calculated with the formula, *w*_pOXA-48+_ / *w*_pOXA-48−_ to correct for the fitness effects of pBCG (see Supplementary Figure 10 for pBGC fitness effects), and the error propagation method was used to calculate the standard error of the resulting value. Note that the fitness effects of pBGC did not correlate with those form pOXA-48 (Pearson’s correlation, R= 0.11, t= 0.66, df= 39, *P*= 0.51). For the 8 strains where pBGC plasmid could not be introduced (Ec13, Kpn10, Kpn11, Kpn19-Kpn23), pOXA-48-carrying and pOXA-48-free isolates were competed against a pBGC-carrying *E. coli* J53^70^ (a sodium azide resistant laboratory mutant of *E. coli* K-12), following the same protocol described above. In general, we prefer to perform competitions assays between isogenic bacteria to avoid interactions between clones that may affect the outcome of the competition for reasons beyond the presence of the plasmid under study (such as bacteriocin production). However, we did not observe any evidence of growth inhibition between the 8 wild type isolates and *E. coli* J53 in the flow cytometry data, and the relative fitness results obtained with these competitions were comparable to those obtained in the isogenic competitions (two-tailed t-test, t= 1.64, df= 11.2, P= 0.13). To confirm that the isogenic competitions and those against *E. coli* J53/pBGC produced similar results, we selected 10 random isolates from the 42 isolates with fitness data calculated from isogenic competitions, and repeated their competitions against *E. coli* J53/pBGC (Supplementary Figure 11). Results showed that relative fitness values calculated with isogenic competitions and those using *E. coli* J53/pBGC presented a good correlation (Pearson’s correlation, R= 0.81, t= 3.96, df= 8, P= 0.004, Supplementary Figure 11). Finally, we performed controls to test for the potential conjugative transfer of pOXA-48 during head-to-head competitions by plating the final time points of the competition assays on amoxicillin-clavulanic acid (with the adequate concentration for each isolate), and chloramphenicol (30 μg/ml). No transconjugants were detected in these controls, showing that the low initial inoculum size we used in the competitions (10,000-fold dilution), and the vigorous shaking of the liquid cultures prevented pOXA-48 conjugation.

### DNA extraction and genome sequencing

Genomic DNA of all the pOXA-48 bearing strains was isolated using the Wizard genomic DNA purification kit (Promega, WI, USA), and quantified using the QuantiFluor dsDNA system (Promega, WI, USA), following manufacturers’ instructions. Whole genome sequencing was conducted at the Wellcome Trust Centre for Human Genetics (Oxford, UK), using the Illumina HiSeq4000 platform with 125 base pair (bp) paired-end reads and at MicrobesNG (Birmingham, UK), using Illumina platforms (MiSeq or HiSeq2500) with 250 bp paired-end reads.

### Bioinformatic analyses

The Illumina sequence reads were trimmed using the Trimmomatic v0.33 tool^71^. SPAdes v3.9.0^72^ was used to generate *de novo* assemblies from the trimmed sequence reads with the –cov-cutoff flag set to ‘auto’. QUAST v4.6.0^73^ was used to generate assembly statistics. Three genomes were dropped from the analysis because of the poor quality of the sequences (2 *E. coli* [Ec09, Ec17] and 1 *K. pneumoniae* [Kpn05]). All the *de novo* assemblies used reached enough quality including total size of 5–7 Mb, and the total number of contigs over 1 kb was lower than 200. Prokka v1.5^74^ was used to annotate the *de novo* assemblies with predicted genes. The seven-gene ST of all the isolates was determined using the multilocus sequence typing (MLST) tool (https://github.com/tseemann/mlst). The plasmid content of each genome was characterised using PlasmidFinder 2.1^75^, and the antibiotic resistance gene content was characterised with ResFinder 3.2^76^ (Supplementary Table 1).

In order to confirm the presence of the entire pOXA-48_K8 plasmid, the sequences belonging to pOXA-48 plasmid in the transconjugants were mapped using as reference the complete sequence of plasmid from the donor strain, which had been previously sequenced by PacBio^31^ (from *K. pneumoniae* k8 – GenBank Accession Number MT441554). Snippy v3.1 (https://github.com/tseemann/snippy) was used to check that no SNPs or indels accumulated in pOXA-48_K8 during strain construction. Coding sequences in pOXA-48 were predicted and annotated using Prokka 1.14.6 software^74^. Plasmid annotation was complemented with the National Center for Biotechnology Information (NCBI) Prokaryotic Genome Annotation Pipeline^77^.

To determine distances between genomes we used Mash v2.0^78^ with the raw sequence reads, and a phylogeny was constructed with mashtree v0.33^79^. For the analysis of the core genome we calculated the genetic relatedness of isolates belonging to *Klebsiella* spp. and to *E. coli* by reconstructing their core genome phylogeny with an alignment of the SNPs obtained with Snippy v3.1 (https://github.com/tseemann/snippy). A maximum-likelihood tree was generated using IQ-TREE with automated detection of the best evolutionary model^80^. The tree was represented with midpoint root using the phylotools package in R (https://github.com/helixcn/phylotools) and visualised using the iTOL tool^81^. We also constructed a distance matrix of the accessory gene network to analyse the accessory genome. To this end, we used AccNET, a tool that allows to infer the accessory genome from the proteomes and cluster them based on protein similarity^46^. The set of representative proteins was used to build a binary matrix (presence/absence of proteins in the accessory genome) in the R-environment and a cladogram to classify the strains according to the accessory genomes. The Euclidean distance was calculated by the ‘dist’ function and a hierarchical clustering was performed with UPGMA using the ‘hclust’ function in the R environment. This cladogram was represented with midpoint root using the phylotools package in R (https://github.com/helixcn/phylotools) and visualised using the iTOL tool^81^.

### Analysis of plasmid fitness effects across bacterial phylogeny

We tested for the presence of phylogenetic signal in core and accessory genomes of *E. coli* and *K. pneumoniae* using several statistical tests available in the *phylosignal* R package^47^. In essence, these analyses are designed to identify statistical dependence between a given continuous trait (relative fitness) and the phylogenetic tree of the taxa from which the trait is measured. Therefore, a positive phylogenetic signal indicates that there is a tendency for related taxa to resemble each other^82^. Several indices have been proposed to identify phylogenetic signal, but the choice among them is not straightforward^83^. We first assayed the methods implemented in the *phyloSignal* function, which produce global measures of phylogenetic signal *(i.e*. across the whole phylogeny). The methods employed were Abouheif’s *C*_mean_, Moran’s I index, Bloomberg’s K and K*, and Pagel’s λ^47^. All methods except Pagel’s λ detected a marginally significant phylogentic signal in the *K. pneumoniae* core genome (Supplementary table 3 [first tab]; 0.11>P>0.02). Abouheif’s *C*_mean_ and Moran’s I (but not Bloomberg’s K and K*, and Pagel’s λ) also detected a marginally significant signal in the *K. pneumoniae* accessory genome tree (Supplementary table 3 [first tab]; P<0.056 for both cases). Intrigued by these results, we used the Local Indicator of Phylogenetic Association (LIPA) based on local Moran’s I, which is meant to detect local hotspots of phylogenetic signal^47,48^. LIPA, implemented in the *lipaMoran* function, computes local Moran’s I indexes for each tip of the phylogeny and a non-parametric test to ascertain statistical significance (Supplementary Figure 4 and Supplementary table 3 [second tab]).

### Plasmid population dynamics model

We used a simple mathematical model of microbial growth under resource limitation to study the role of the DFE in the ecological dynamics of a plasmid spreading in a bacterial population^14^. Bacterial growth rate was modelled as a saturating function of the environmental resource concentration, *R*,

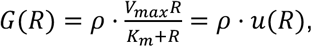

where *ρ* denotes the cell’s efficiency to convert resource molecules into biomass and *u*(*R*) a resource uptake function that depends on the maximum uptake rate *(V_max_*) and a half-saturation constant *(K_m_*). If we denote with *B_p_* the density of plasmid-bearing cells and with *B*_0_ the density of plasmid-free cells (each with its own growth kinetic parameters and growth functions denoted *G_p_*(*R*) and *G*_0_(*R*), respectively), then the density of each subpopulation can be described by a system of ordinary differential equations:

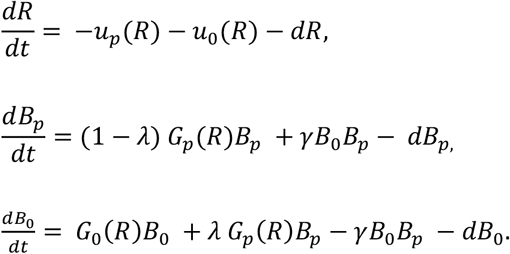

where *λ* represents the rate of segregational loss rate and *d* a dilution parameter. Moreover, we represent with *γ* the rate of conjugative transfer, and therefore we model plasmid conjugation as a function of the densities of donor and recipient cells. By numerically solving the system of equations (using standard differential equations solvers in Matlab), we obtain the final density of each bacterial type in an experiment of *T* = 24 units of time with *d* = 0 (to replicate the batch culture conditions used to estimate the DFE experimentally).

Growth kinetic parameters were determined with a Markov chain Monte Carlo method (MCMC; scripts coded in R and available at http://www.github.com/esb-lab/pNUK73/) applied to growth curves of each strain growing in isolation, with and without plasmids. This data fitting algorithm implements a Metropolis-Hastings sampler with a burn-in parameter of 0.2 and executed for 5 × 10^6^ iterations, or until achieving convergence of the Markov chains (see Supplementary Figure 5 for an example and Supplementary Table 5 for parameters values estimated for each strain).

### Stochastic simulations of polymicrobial communities

Numerical experiments were performed by randomly sampling *N* = 1× 10^4^ cells from the parameter distribution obtained after applying the MCMC algorithm to all 50 strains and fitting a bivariate Normal distribution. We then assembled 5,000 synthetic communities composed of a random subset of M < N different strains sampled from this distribution, and solved a multi-strain extension of the population dynamics model. For each numerical experiment, the total density of strain *i* would be 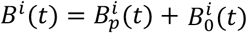 where 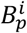 and 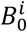 denote, respectively, the densities of plasmid-bearing and plasmid-free cells of type 1 ≤ *i ≤ M*. To model the fitness effects of bearing plasmids, we introduced a parameter, *σ*, such that when *σ* = 0, the fitness difference between 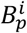 and 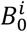 corresponds to a fixed reduction in growth rate (corresponding to a DFE with variance 0 and mean *w* = 0.985). Conversely, if *σ >* 0, then growth kinetic parameters for each plasmid-bearing strain in the community were determined by sampling *s_i_* from a Normal distribution, *N*(0, *σ*^2^), and multiplying both 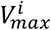 and *ρ^i^* by a factor of (1 + *s_i_*).

As with the single-strain model, we consider segregational loss as a transition from 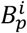 to 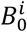 occurring at a rate *λ*, but now we also consider that plasmid-free cells can acquire plasmids via conjugation from any plasmid-bearing strain in the community, at a constant rate *γ*, and with equal probability of transferring between different bacterial hosts. Therefore, we obtain a system of 2*M* + 1 differential equations that can be written, for each strain *i*, as follows:

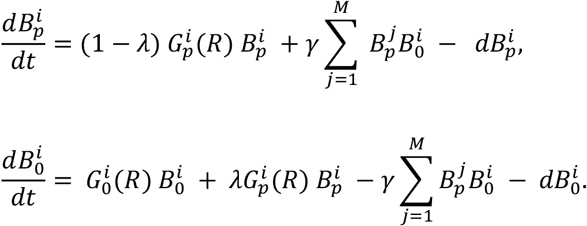

Furthermore, if 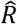 represents the input of resource into the system, then

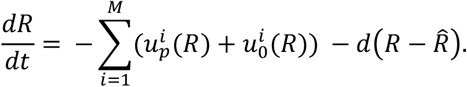

Initial bacterial densities were determined by first running the system forward (with all strains initially present at equal densities) for *T* = 24 time units, and then clearing all plasmid-free cells from the population. This assumption is consistent with patients receiving antimicrobial therapy that clears all susceptible (plasmid-free) cells from the microbiota or, in an experimental microcosm, to a round of growth in selective media after an overnight culture. As we are interested in the long-term population dynamics, we ran each simulation starting from the aforementioned initial condition until the plasmid fraction was below a threshold *ϵ* > 0 (i.e. plasmid extinction), the plasmid fraction was near 100% and the total plasmid-free density was below *ϵ* (i.e. plasmid fixation), or wild-type and transconjugant sub-populations appeared to co-exist indefinitely in the population (either in equilibrium or exhibiting oscillatory behaviour, as illustrated in Supplementary Figures 7 and 8).

### Statistical analyses

The statistical tests used are indicated in the text. Analyses were performed using R (v. 3.5.0).

## Acknowledgements

This work was supported by the European Research Council under the European Union’s Horizon 2020 research and innovation programme (ERC grant agreement no. 757440-PLASREVOLUTION) and by the *Instituto de Salud Carlos III* (co-funded by European Development Regional Fund “a way to achieve Europe”) grant PI16-00860. RC acknowledges financial support from European Commission (grant R-GNOSIS-FP7-HEALTH-F3-2011-282512) and *Plan Nacional de I+D+i2013–2016* and *Instituto de Salud Carlos III, Subdirección General de Redes y Centros de Investigación Cooperativa, Ministerio de Economía, Industria y Competitividad*, Spanish Network for Research in Infectious Diseases (REIPIR D16/0016/0011) co-financed by European Development Regional Fund “A way to achieve Europe” (ERDF), Operative program Intelligent Growth 2014–2020. ASM is supported by a Miguel Servet Fellowship (MS15-00012). JRB is a recipient of a Juan de la Cierva-Incorporación Fellowship (IJC2018-035146-I) co-funded by *Agencia Estatal de Investigación del Ministerio de Ciencia e Innovación*. MH-G was supported with a contract from *Instituto de Salud Carlos III*, Spain (iP-FIS program, ref. IFI14/00022). RPM was supported by PAPIIT-UNAM (IN209419) and CONACYT (Ciencia Básica grant A1-S-32164). We thank the Oxford Genomics Centre at the Wellcome Centre for Human Genetics (funded by Wellcome Trust grant reference 203141/Z/16/Z) for the generation and initial processing of the sequencing data.

## Author Contributions

ASM, AAdV and RPM conceived the study. RC designed and supervised sampling and collection of R-GNOSIS bacterial isolates. MHG, PRG collected the bacterial isolates and performed bacterial characterization. AAdV performed the experimental work with help from JRB and JdLF. AAdV, JRB and JdLF analysed experimental results. RLS and JRB performed the bioinformatic/phylogenetic analyses. RPM developed the mathematical model and computer simulations. ASM supervised the study. ASM, AAdV and RPM wrote the initial draft of the manuscript and all the authors contributed to the final version of the manuscript and approved it.

## Data availability

The sequences generated and analysed during the current study are available in the Sequence Read Archive (SRA), BioProject ID: PRJNA641166, https://www.ncbi.nlm.nih.gov/sra/PRJNA641166.

## Code availability

The code generated during the current study is available in GitHub, http://www.github.com/esb-lab/pOXA48/

## Supplementary Information

**Supplementary Figure 1.**
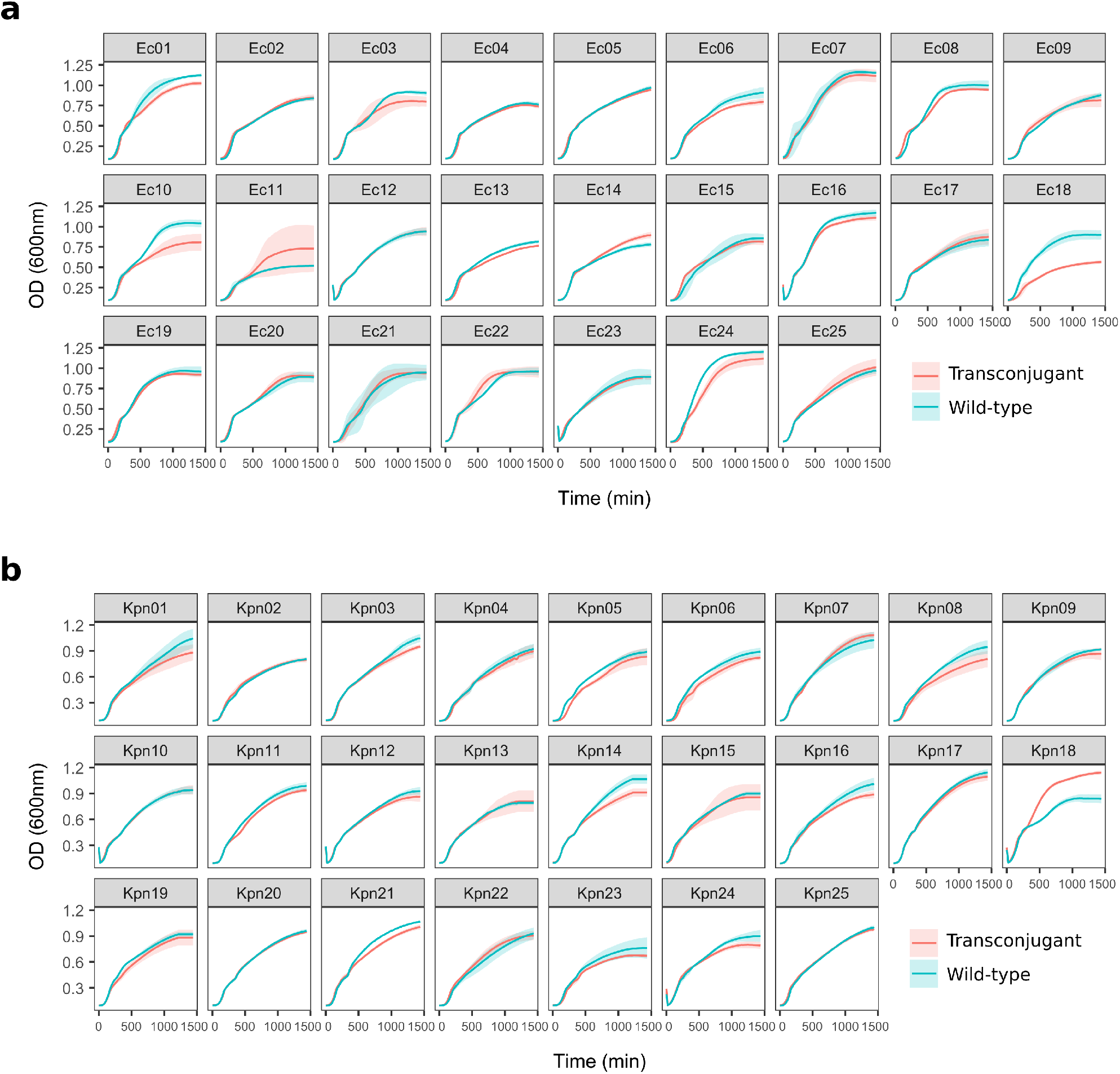
Growth curves of wild-type isolates and pOXA-48-carrying transconjugants. Growth curves of pOXA48-free (wild-type, blue) and pOXA-48-carrying (transconjugant, red) for every (a) *E. coli* and (b) *Klebsiella* spp. analysed in this study. The lines represent the average of five biological replicates and the shaded area indicates 95% confidence intervals.

**Supplementary Figure 2.**
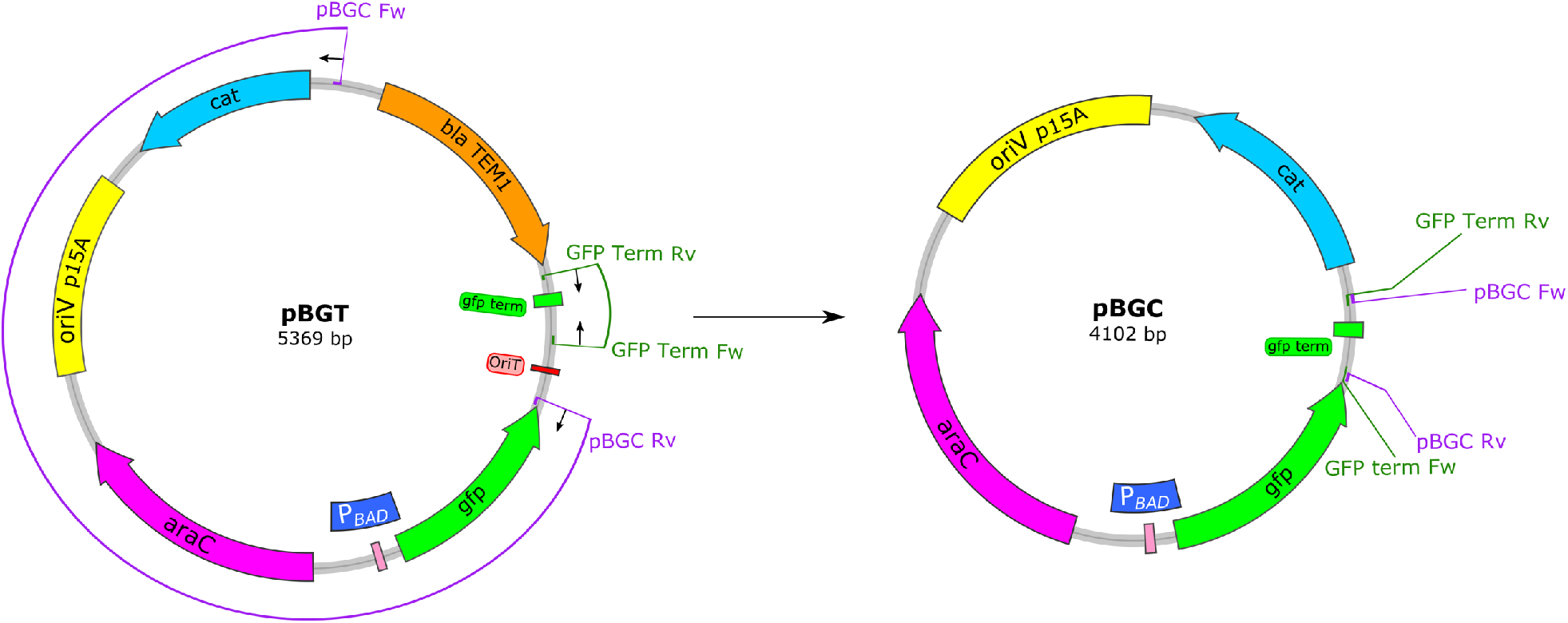
Construction of plasmid pBGC. Schematic representation of the construction of plasmid pBGC (accession number MT702881) form plasmid pBGT^68^. Two segments of pBGT were amplified using primers with added cohesive ends (pBGC Fw/Rv and GFP Term Fw/Rv, Supplementary Table 4). pBGC plasmid resulted from the Gibson assembly of the amplified fragments. The reading frames for genes are shown as arrows, with the direction of transcription indicated by the arrowhead. The origin of replication (oriV), origin of transfer (oriT), and *P_BAD_* promoter are also indicated.

**Supplementary Figure 3.**
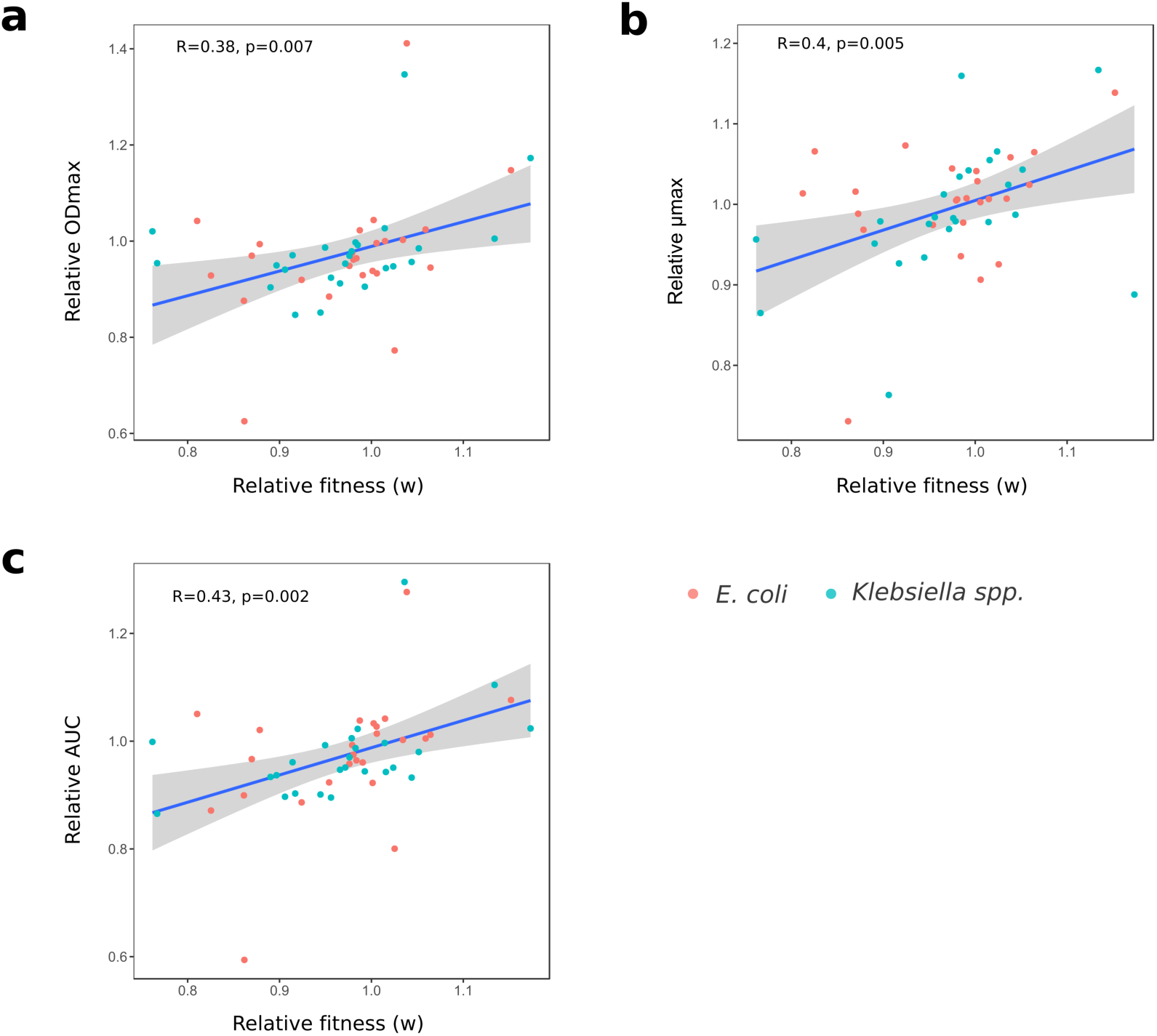
Correlation between relative growth curve parameters and relative fitness. Correlation between relative growth curve parameters (a) maximum optical density (OD_max_), (b) maximum growth rate (μ_max_), and (c) area under the growth curve (AUC), and relative fitness values obtained from competition assays for each strain. The blue line represents the linear regression model and the grey shading represents 95% confidence intervals. Points represent each relative value (red, *E*. coli and blue, *Klebsiella* spp.). Pearson’s correlation (R) and p-value are indicated. As expected, maximum optical density, maximum growth rate and area under the growth curve are positively correlated with relative fitness.

**Supplementary Figure 4.**
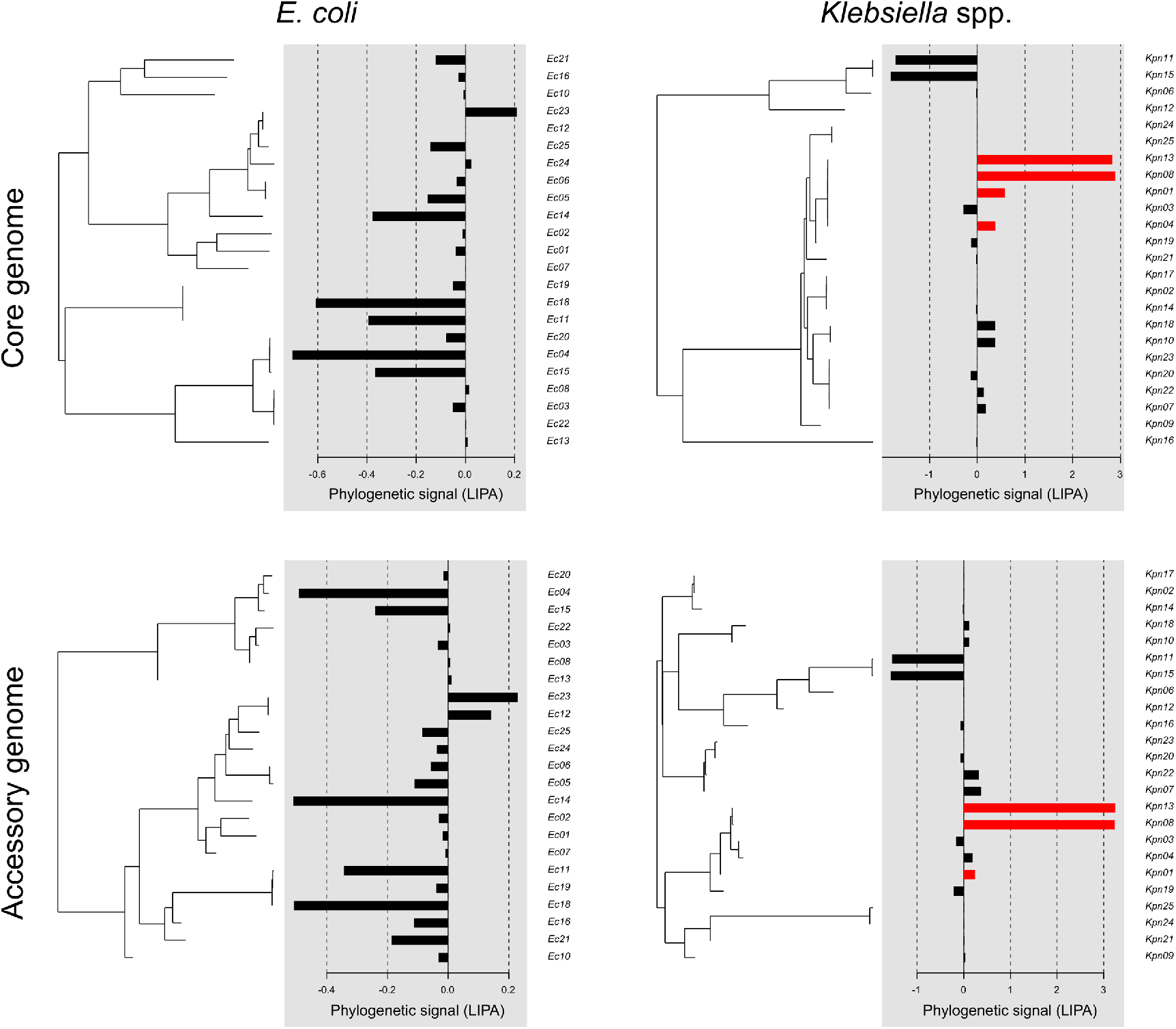
Local Indicator of Phylogenetic Association (LIPA) analyses. Phylogenetic trees for core (upper panels) and accessory (lower panels) genomes obtained for *E. coli* (left) and *Klebsiella* spp. (right). Bar plots show the LIPA score associated with each tip of the phylogeny, with higher values representing a stronger phylogenetic signal. Red colour indicates statistically significant LIPA scores (*i.e*. phylogenetic signal, Supplementary Table 3).

**Supplementary Figure 5.**
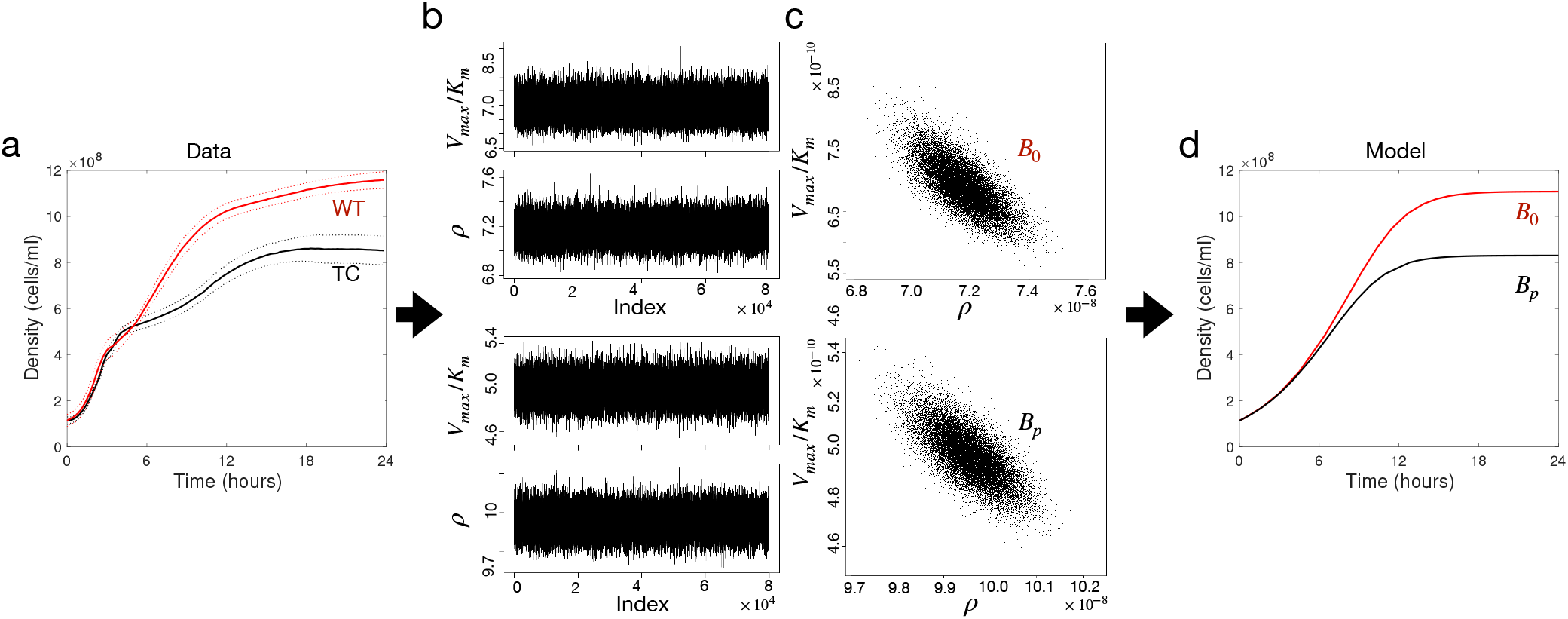
Model parametrization. (a) Bacterial density of strain Kpn18 as a function of time obtained from the optical density of wild-type (red) and transconjugant (black) strains growing in isolation. (b) Traces of chains for parameters *V_max_/K_m_* (above) and *ρ* (below) obtained by fitting a simple Monod model to growth curve data using a Metropolis-Hastings Markov chain Monte Carlo method (MCMC). (c) 2-dimensional posterior distributions obtained for each strain (top: *B*_0_, bottom: *B_p_*). (d) Numerical solutions of the model using parameters selected randomly from the posterior distribution and with initial conditions determined from the experimental growth curves.

**Supplementary Figure 6.**
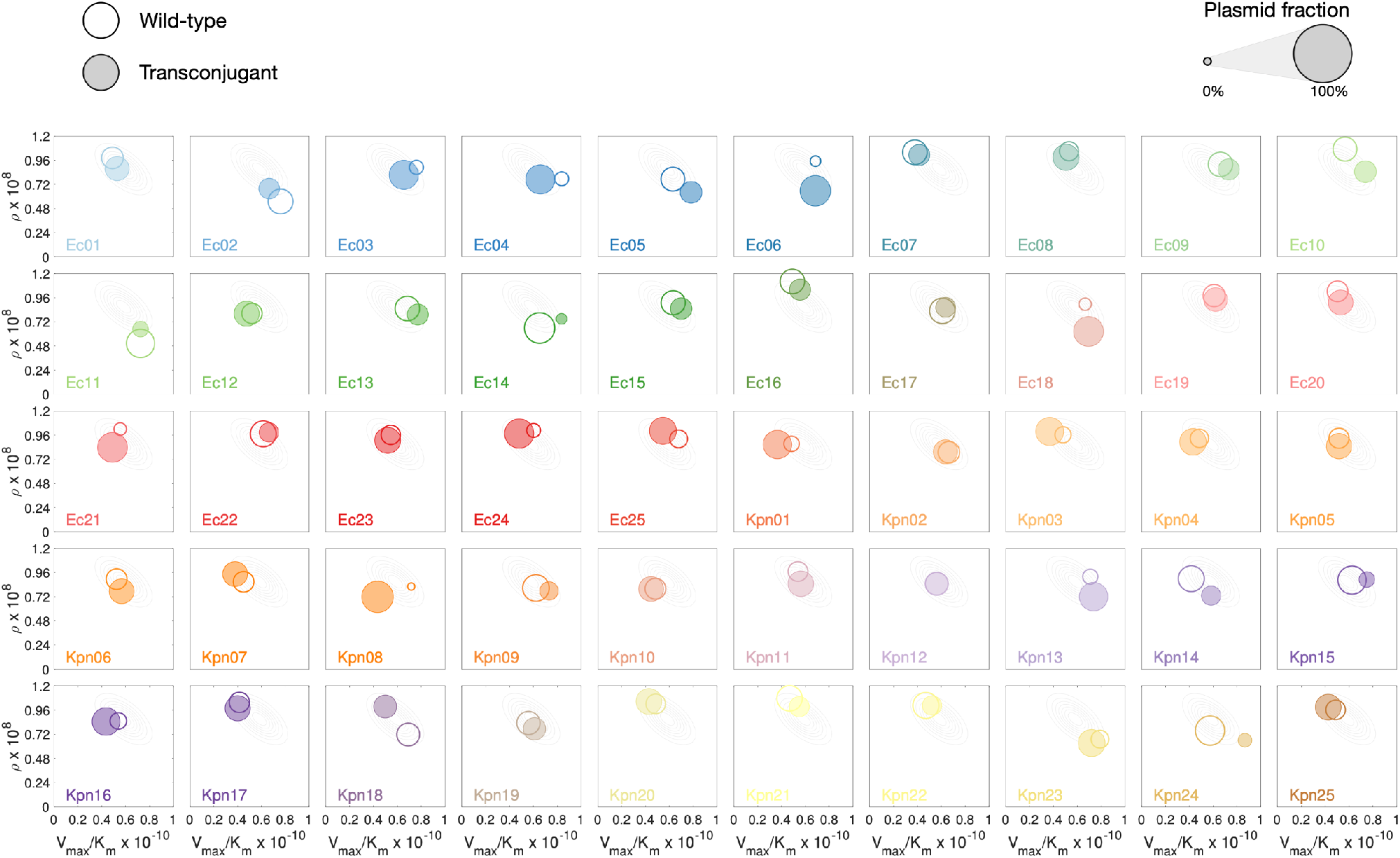
*In silico* competition experiments. Each box represents a theoretical pair-wise competition experiment between plasmid-free (open circles) and plasmid-bearing cells (filled circles). The diameter of each circle is proportional to the relative fraction of the population, a value estimated by numerically solving the model for *T* = 24 with parameter values obtained from the posterior distribution of each strain. Horizontal axis represents the specific affinity *(V_max_/K_m_*) and the vertical axis the cell’s resource conversion rate (*ρ*).

**Supplementary Figure 7.**
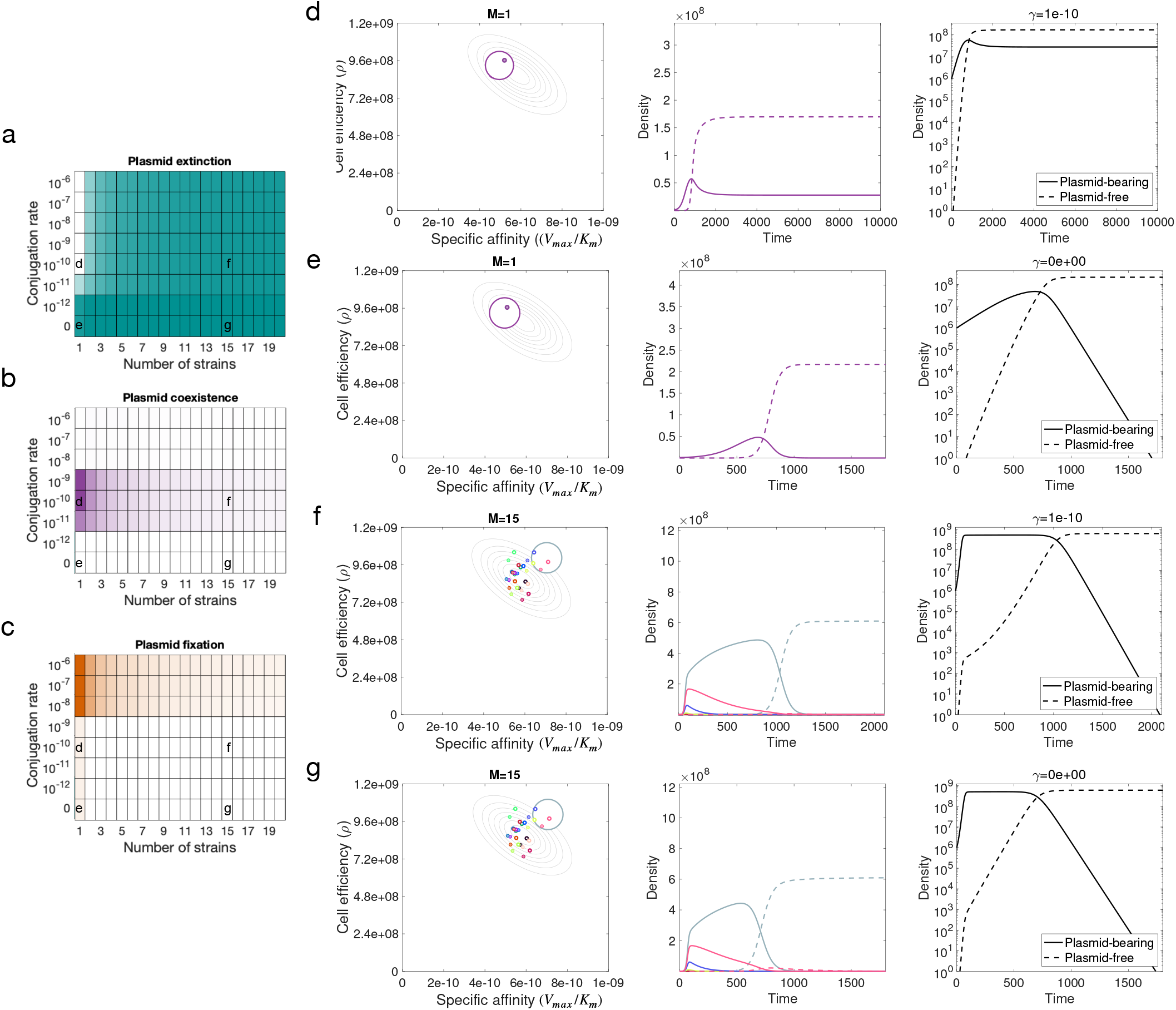
Effect of conjugation and community complexity in plasmid population dynamics in the absence of a DFE. Numerical simulations of the population dynamics model performed over a range of conjugation rates and number of strains in the community. In this case, all strains exhibit a reduction of fitness when carrying the plasmid (a DFE with mean w= 0.985 and variance 0). The colour of each box in the grid corresponds to the percentage of 5,000 random communities that exhibited: a) plasmid extinction (total plasmid frequency was below a threshold), b) plasmid-bearing and plasmid-free cells co-exist in the population, and c) every cell in the population carries the plasmid at the end of the experiment. (d-g) Example of relative abundances over time for a range of conjugation rates in a community composed of 1 (d,e) and 15 (f,g) strains, with segregation rate *λ* = 1 × 10^−8^ and conjugation rate *γ* = 10^−10^ (d,f) or *γ* = 0 (e,g). The left-hand column illustrates the growth kinetic parameters for each strain (empty circles denote plasmid-free cells and filled circles plasmid-bearing cells, with diameters proportional to their final relative abundances). Middle column shows the density of each subpopulation as a function of time (dotted lines denote plasmid-free strains and solid lines subpopulations carrying the plasmid). Right-hand column shows semilog plots with the total fraction of cells with and without plasmids (solid and dotted lines, respectively). As plasmid-bearing is associated with a fitness cost, then the plasmid is only maintained in the population at high conjugation rates.

**Supplementary Figure 8.**
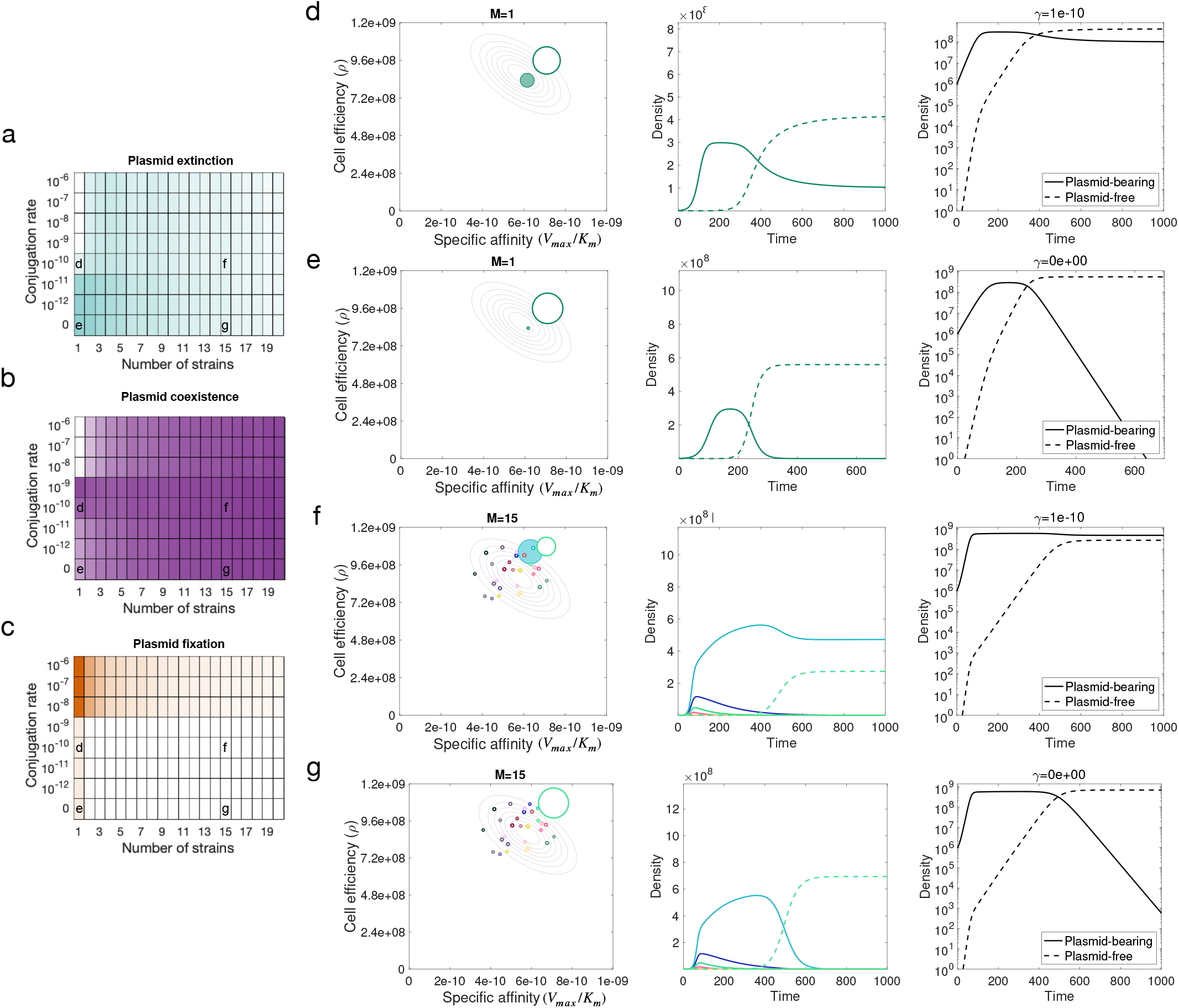
Effect of conjugation and community complexity in plasmid population dynamics in the presence of a DFE. Numerical simulations of the population dynamics model performed over a range of conjugation rates and number of strains in the community. This case corresponds to a wide DFE (mean w = 0.985 and variance 0.007). The colour of each box in the grid corresponds to the percentage of 5,000 random communities that exhibited: a) plasmid extinction (total plasmid frequency was below a threshold), b) plasmid-bearing and plasmid-free cells co-exist in the population, and c) every cell in the population carries the plasmid at the end of the experiment. d-g) Example of relative abundances over time for a range of conjugation rates in a community composed of 1 (d,e) and 15 (f,g) strains, with segregation rate *λ* = 1 × 10^−8^ and conjugation rate *γ* = 10^−10^ (d,f) or *γ* = 0 (e,g). The left-hand column illustrates the growth kinetic parameters for each strain (empty circles denote plasmid-free cells and filled circles plasmid-bearing cells, with diameters proportional to their final relative abundances). Middle column shows the density of each subpopulation as a function of time (dotted lines denote plasmid-free strains and solid lines subpopulations carrying the plasmid). Right-hand column shows semilog plots with the total fraction of cells with and without plasmids (solid and dotted lines, respectively). Note how a wide DFE allows plasmids to persist, even at very low conjugation rates.

**Supplementary Figure 9.**
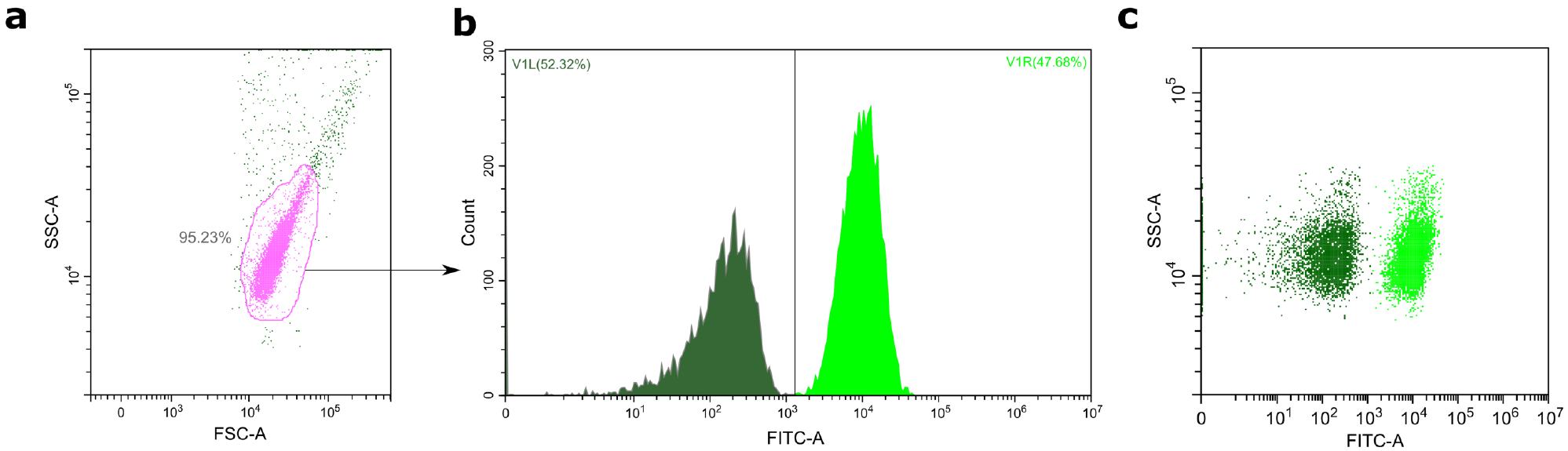
Determination of different cells types in competition assays using flow cytometry. We used flow cytometry to differentiate between GFP-producing and -non-producing cells. (a) we used forward versus side scatter (FSC vs SSC) gating to identify bacterial cells in the sample. (b-c) GFP-producing (bright green) and -non-producing (dark green) cells were differentiated using the FITC-A (fluorescein isothiocyanate) channel, allowing us to measure the proportion of each competitor in the mix.

**Supplementary Figure 10.**
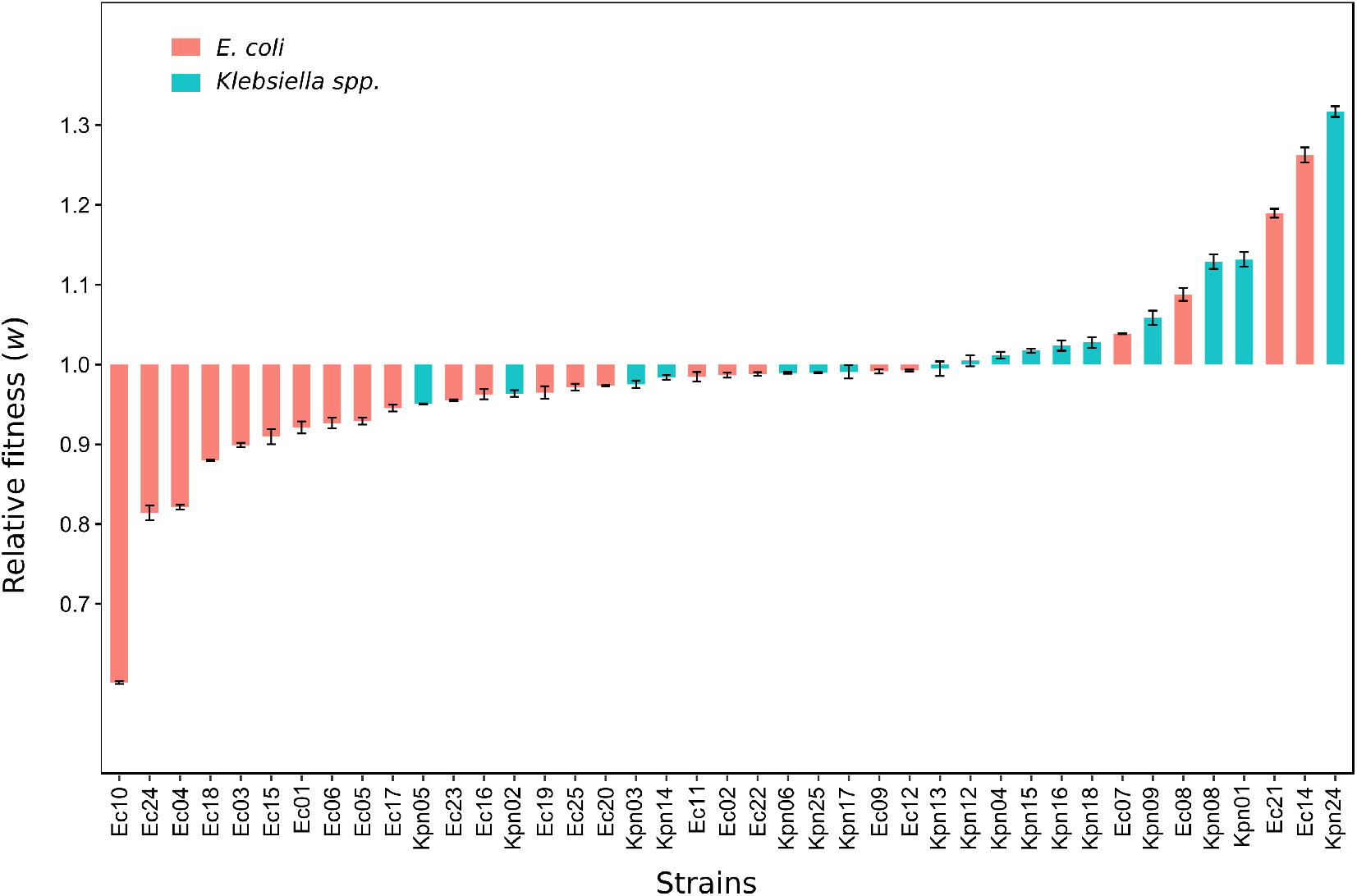
Distribution of pBGC fitness effects. Relative fitness (*w*) of pBGC-carrying clones compared to plasmid-free clones, obtained from competition assays (red, *E. coli* and blue, *Klebsiella* spp.). Values below 1 indicate a reduction in *w* and values above 1 indicate an increase in *w* due to pBGC acquisition. Bars represent the average of five independent experiments and error bars represent the standard error of the mean. Note that the fitness effects of pBGC did not correlate with those form pOXA-48 (Pearson’s correlation, R= 0.11, t= 0.66, df= 39, *P*= 0.51).

**Supplementary Figure 11.**
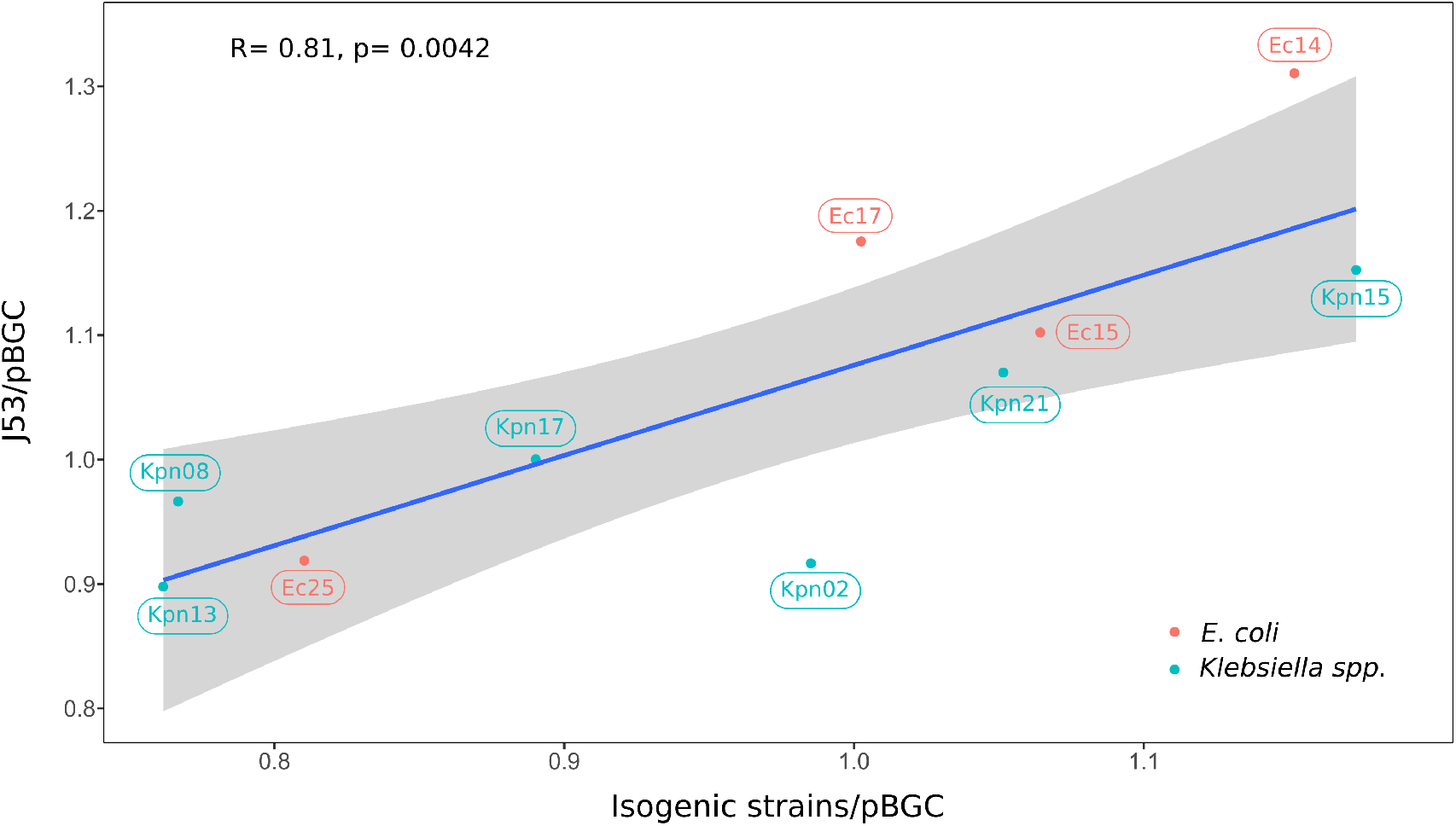
Correlation between relative fitness values calculated in competitions vs. *E. coli* J53/pBGC or isogenic clones with pBGC. Correlation between relative fitness values obtained from competitions assays using pBGC-carrying isogenic isolates and pBGC-carrying *E*. coli J53 for ten different isolates. The blue line represents the linear regression model and the grey shading represents 95% confidence intervals. Blue points correspond to *Klebsiella* spp. isolates and red points to *E. coli* isolates. Labels indicate isolates names. Pearson’s correlation (R) and p-value are indicated.

**Supplementary Table 4.**
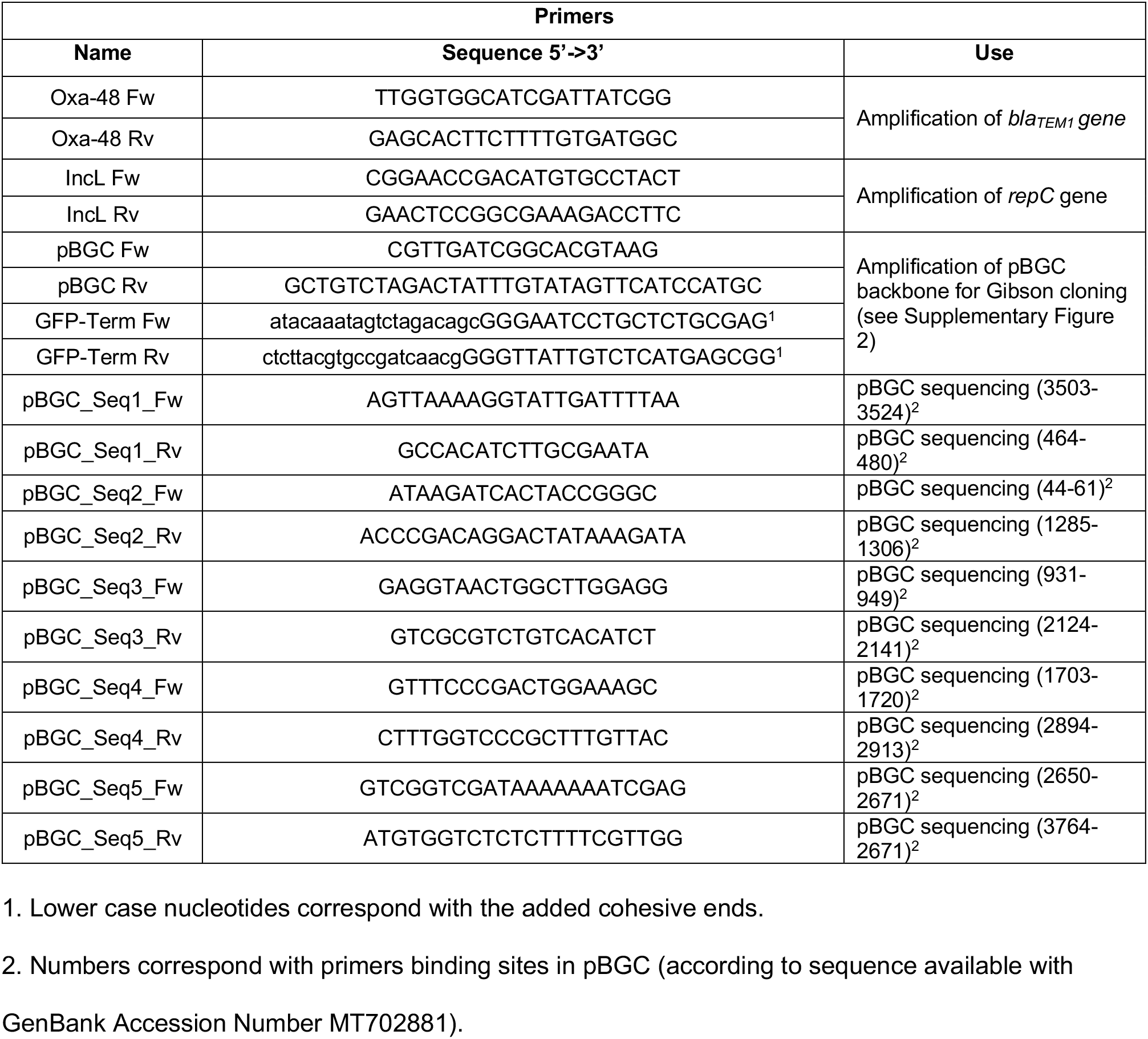
Primers used in this study.

**Supplementary Table 5.** Growth kinetic parameters of each strain obtained using the MCMC algorithm.

| Strain | Wild-type |  | Transconjugant |  |
| --- | --- | --- | --- | --- |
| | $V_{max}/K$ | $\rho$ | $V_{max}/K$ | $\rho$ |
| Ec01 | $4.895 \times 10^{-10}$ | $9.844 \times 10^8$ | $5.283 \times 10^{-10}$ | $8.776 \times 10^8$ |
| Ec02 | $7.603 \times 10^{-10}$ | $5.504 \times 10^8$ | $6.648 \times 10^{-10}$ | $6.786 \times 10^8$ |
| Ec03 | $7.597 \times 10^{-10}$ | $8.900 \times 10^8$ | $6.538 \times 10^{-10}$ | $8.151 \times 10^8$ |
| Ec04 | $8.373 \times 10^{-10}$ | $7.764 \times 10^8$ | $6.576 \times 10^{-10}$ | $7.702 \times 10^8$ |
| Ec05 | $6.299 \times 10^{-10}$ | $7.727 \times 10^8$ | $7.831 \times 10^{-10}$ | $6.416 \times 10^8$ |
| Ec06 | $6.852 \times 10^{-10}$ | $9.518 \times 10^8$ | $6.839 \times 10^{-10}$ | $6.567 \times 10^8$ |
| Ec07 | $3.786 \times 10^{-10}$ | $1.040 \times 10^9$ | $4.154 \times 10^{-10}$ | $1.013 \times 10^9$ |
| Ec08 | $5.304 \times 10^{-10}$ | $1.050 \times 10^9$ | $5.053 \times 10^{-10}$ | $9.900 \times 10^8$ |
| Ec09 | $6.610 \times 10^{-10}$ | $9.195 \times 10^8$ | $7.311 \times 10^{-10}$ | $8.694 \times 10^8$ |
| Ec10 | $5.660 \times 10^{-10}$ | $1.071 \times 10^9$ | $7.368 \times 10^{-10}$ | $8.472 \times 10^8$ |
| Ec11 | $7.248 \times 10^{-10}$ | $5.064 \times 10^8$ | $7.248 \times 10^{-10}$ | $6.483 \times 10^8$ |
| Ec12 | $5.200 \times 10^{-10}$ | $8.042 \times 10^8$ | $4.752 \times 10^{-10}$ | $8.016 \times 10^8$ |
| Ec13 | $6.831 \times 10^{-10}$ | $8.528 \times 10^8$ | $7.711 \times 10^{-10}$ | $7.932 \times 10^8$ |
| Ec14 | $6.515 \times 10^{-10}$ | $6.589 \times 10^8$ | $8.356 \times 10^{-10}$ | $7.500 \times 10^8$ |
| Ec15 | $6.322 \times 10^{-10}$ | $9.106 \times 10^8$ | $6.998 \times 10^{-10}$ | $8.528 \times 10^8$ |
| Ec16 | $4.908 \times 10^{-10}$ | $1.122 \times 10^9$ | $5.550 \times 10^{-10}$ | $1.040 \times 10^9$ |
| Ec17 | $6.089 \times 10^{-10}$ | $8.298 \times 10^8$ | $6.366 \times 10^{-10}$ | $8.647 \times 10^8$ |
| Ec18 | $6.655 \times 10^{-10}$ | $8.963 \times 10^8$ | $6.939 \times 10^{-10}$ | $6.250 \times 10^8$ |
| Ec19 | $6.080 \times 10^{-10}$ | $9.812 \times 10^8$ | $6.200 \times 10^{-10}$ | $9.424 \times 10^8$ |
| Ec20 | $5.033 \times 10^{-10}$ | $1.024 \times 10^9$ | $5.296 \times 10^{-10}$ | $9.133 \times 10^8$ |
| Ec21 | $5.546 \times 10^{-10}$ | $1.021 \times 10^9$ | $4.897 \times 10^{-10}$ | $8.386 \times 10^8$ |
| Ec22 | $6.148 \times 10^{-10}$ | $9.736 \times 10^8$ | $6.619 \times 10^{-10}$ | $9.886 \times 10^8$ |
| Ec23 | $5.443 \times 10^{-10}$ | $9.635 \times 10^8$ | $5.186 \times 10^{-10}$ | $9.094 \times 10^8$ |
| Ec24 | $6.010 \times 10^{-10}$ | $1.008 \times 10^9$ | $4.808 \times 10^{-10}$ | $9.768 \times 10^8$ |
| Ec25 | $6.784 \times 10^{-10}$ | $9.237 \times 10^8$ | $5.465 \times 10^{-10}$ | $1.004 \times 10^9$ |
| Kpn01 | $4.841 \times 10^{-10}$ | $8.746 \times 10^8$ | $3.652 \times 10^{-10}$ | $8.673 \times 10^8$ |
| Kpn02 | $6.638 \times 10^{-10}$ | $7.905 \times 10^8$ | $6.352 \times 10^{-10}$ | $7.952 \times 10^8$ |
| Kpn03 | $4.802 \times 10^{-10}$ | $9.639 \times 10^8$ | $3.686 \times 10^{-10}$ | $9.975 \times 10^8$ |
| Kpn04 | $4.844 \times 10^{-10}$ | $9.288 \times 10^8$ | $4.303 \times 10^{-10}$ | $8.936 \times 10^8$ |
| Kpn05 | $5.151 \times 10^{-10}$ | $9.303 \times 10^8$ | $5.155 \times 10^{-10}$ | $8.513 \times 10^8$ |
| Kpn06 | $5.240 \times 10^{-10}$ | $8.952 \times 10^8$ | $5.664 \times 10^{-10}$ | $7.742 \times 10^8$ |
| Kpn07 | $4.498 \times 10^{-10}$ | $8.682 \times 10^8$ | $3.785 \times 10^{-10}$ | $9.458 \times 10^8$ |
| Kpn08 | $7.163 \times 10^{-10}$ | $8.211 \times 10^8$ | $4.334 \times 10^{-10}$ | $7.188 \times 10^8$ |
| Kpn09 | $6.207 \times 10^{-10}$ | $8.064 \times 10^8$ | $7.302 \times 10^{-10}$ | $7.772 \times 10^8$ |
| Kpn10 | $4.833 \times 10^{-10}$ | $7.990 \times 10^8$ | $4.467 \times 10^{-10}$ | $7.967 \times 10^8$ |
| Kpn11 | $5.380 \times 10^{-10}$ | $9.710 \times 10^8$ | $5.620 \times 10^{-10}$ | $8.490 \times 10^8$ |
| Kpn12 | $5.620 \times 10^{-10}$ | $8.490 \times 10^8$ | $5.620 \times 10^{-10}$ | $8.490 \times 10^8$ |
| Kpn13 | $7.086 \times 10^{-10}$ | $9.195 \times 10^8$ | $7.364 \times 10^{-10}$ | $7.206 \times 10^8$ |
| Kpn14 | $4.162 \times 10^{-10}$ | $8.994 \times 10^8$ | $5.828 \times 10^{-10}$ | $7.306 \times 10^8$ |
| Kpn15 | $6.243 \times 10^{-10}$ | $8.854 \times 10^8$ | $7.489 \times 10^{-10}$ | $8.917 \times 10^8$ |
| Kpn16 | $5.386 \times 10^{-10}$ | $8.523 \times 10^8$ | $4.359 \times 10^{-10}$ | $8.465 \times 10^8$ |
| Kpn17 | $4.144 \times 10^{-10}$ | $1.037 \times 10^9$ | $4.002 \times 10^{-10}$ | $9.786 \times 10^8$ |
| Kpn18 | $6.906 \times 10^{-10}$ | $7.173 \times 10^8$ | $4.977 \times 10^{-10}$ | $9.954 \times 10^8$ |
| Kpn19 | $5.578 \times 10^{-10}$ | $8.319 \times 10^8$ | $6.083 \times 10^{-10}$ | $7.739 \times 10^8$ |
| Kpn20 | $4.857 \times 10^{-10}$ | $1.022 \times 10^9$ | $4.287 \times 10^{-10}$ | $1.046 \times 10^9$ |
| Kpn21 | $4.674 \times 10^{-10}$ | $1.077 \times 10^9$ | $5.487 \times 10^{-10}$ | $9.948 \times 10^8$ |
| Kpn22 | $4.708 \times 10^{-10}$ | $1.006 \times 10^9$ | $5.224 \times 10^{-10}$ | $1.006 \times 10^9$ |
| Kpn23 | $7.897 \times 10^{-10}$ | $6.709 \times 10^8$ | $7.201 \times 10^{-10}$ | $6.328 \times 10^8$ |
| Kpn24 | $5.736 \times 10^{-10}$ | $7.568 \times 10^8$ | $8.670 \times 10^{-10}$ | $6.601 \times 10^8$ |
| Kpn25 | $4.873 \times 10^{-10}$ | $9.630 \times 10^8$ | $4.255 \times 10^{-10}$ | $9.896 \times 10^8$ |

